# Respiratory pauses highlight sleep architecture in mice

**DOI:** 10.1101/2024.03.27.586921

**Authors:** Giulio Casali, Camille Miermon, Geoffrey Terral, Pascal Ravassard, Tiphaine Dolique, Evan Harrell, Edith Lesburguères, David Jarriault, Frédéric Gambino, Nicolas Chenouard, Lisa Roux

## Abstract

Brain activity and breathing rate influence each other but it remains unclear how fine respiratory features vary across vigilance states. Using simultaneous nasal pressure and hippocampal local field potential recordings in freely-moving mice, we show that the position of respiratory pauses within breathing cycles distinguish Wake, Rapid Eye Movement (REM) and non-REM (NREM) sleep states. Model-based predictions of vigilance states based on respiratory features perform well even on animals outside of the training set, suggesting the rules are generalized. Respiratory features underwent specific changes at state transitions, such as progressive elimination of pauses after inhalation foreshadowing REM. During NREM, respiratory changes predicted moment-to moment sigma power variations beyond movement-defined packets delineated by micro-arousals, as pauses after inhalation fragmented NREM sleep into ∼30s windows of high sigma power. Overall, our findings reveal that respiratory features structure the macro- and micro-architecture of sleep, opening new windows into brain states through respiration.

## Introduction

While respiration is crucial for survival by its peripheral action (1), it also shares strong functional links with neuronal networks (2–5). Changes in brain activity between Wake, Rapid-Eye-Movement (REM) and Non-Rapid-Eye-Movement (NREM) sleep are accompanied by significant modifications in respiratory behavior, both in humans and animal species (6–15). The two main sub-stages of sleep can be identified based on electrophysiological (e.g. local field potential - LFP - or electroencephalogram - EEG) and muscular activity (e.g. electromyogram - EMG): REM sleep is typically associated with desynchronized brain activity, ocular saccades and nearly absent muscular tone (16–19), and NREM sleep with enhanced cortical slow wave activity and higher muscular tone. In NREM sleep, recent works also highlighted the presence of 20 to 120 sec time-windows delimited by micro-arousals that are referred to as “NREM packets” (20–22). These packets presumably generate infra-slow oscillations during NREM sleep both in humans and rodent models (23, 24) and show specific neuromodulatory and LFP spectral characteristics, notably an increased sigma-band (10-20 Hz) power and heightened spindle density in their center (20–22). Spindles consist of transient bursts of sigma oscillations generated by thalamocortical circuits that also influence hippocampal LFP (25) and which are presumed to support memory consolidation during NREM (19, 26, 27). As they constrain spindles occurrence, NREM packets have been shown to orchestrate memory (21) and homeostatic (20) processes in hippocampo-cortical circuits. They also determine sleep fragility, dictating the propensity of mice to wake up in response to external stimuli (21). Because the fine microarchitecture of NREM is central to understanding its role in cognitive functions, and since respiration can shape neuronal activity in sleep (6–8, 11, 13–15), it becomes necessary to determine how respiration accompanies these phenomena.

Previous works showed that respiration rate and amplitude decrease during sleep as compared to Wake (6, 7, 14), with REM showing higher respiration rates as compared to NREM (11, 28), although some discrepancies exist regarding REM (29, 30). However, respiration is a complex phenomenon that can be described by many more features than just rate and amplitude (31, 32). For instance, the precise dynamics and volumes of inhalation and exhalation are important features that also vary across vigilance states (7). In addition, epochs with no airflow nested within breathing cycles have been observed in human (31) and rodent (32) respiration recordings in physiological conditions. These respiratory pauses have so far never been specifically studied and could potentially relates to brain states and transitions between them.

Methodological advances based on pressure sensors that allow a precise monitoring of nasal airflow in rodents (32–36) and the development of new analysis pipelines dedicated to respiratory pressure signals (31) are offering an unprecedented opportunity to explore these questions. By combining nasal pressure monitoring with LFP recordings in the CA1 region of the hippocampus of freely moving mice, our work aimed at understanding how the fine features of respiration – and in particular pauses - co-vary relative to one another and how they behave in different vigilance states. Because respiratory changes around state transitions and NREM packets remain largely uncharacterized, we also asked whether respiratory variations precede or follow transitions undergone by hippocampal networks. Finally, we tested the hypothesis that they can capture the micro-architecture of NREM sleep and its so-called “packets”.

## Results

### Monitoring respiration and brain states in freely-behaving mice highlights changes of respiration kinetics across states

In order to measure breathing in feely-moving mice with high accuracy while minimally interfering with the physiological alternations of vigilance states, we attached light and portable pressure sensors to intranasal cannulas and monitored air pressure in the nasal cavity (33, 34, 37) (**Fig. 1A**; **Methods**). Pressure sensors, unlike thermocouples (**Supplementary Fig. 1A**), are sensitive enough to detect fine changes in nasal airflow including during sleep states (32) (**Fig. 1B, Supplementary Fig. 1A, Supplementary Fig. 2**). We verified that the intranasal cannula implantation did not interfere with basic olfactory capacity (odor detection test**; Supplementary Fig. 1B**) and spontaneous exploratory behavior (**Supplementary Fig. 1B**). As hippocampal LFP is well characterized across brain states (38–40), we implanted 8 mice with intranasal cannulas and silicon probes targeting the CA1 region of the dorsal hippocampus (**Fig. 1A**). We used CA1 LFP in combination with locomotion/accelerometer signals to define the three main brain states (**Methods**): Wake (high locomotion and high theta/delta power ratio), NREM sleep (immobility and low theta/delta power ratio), and REM sleep (immobility with prominent theta-band increase). The freely-moving conditions allowed us to monitor brain states and respiration throughout long (> 2h) periods of time as mice spontaneously alternated between these three brain states (mean ± SEM; Wake, 9090 ± 506 seconds; REM, 466 ± 47 seconds; NREM, 6368 ± 400 seconds; N = 38 sessions).

**Figure 1.**
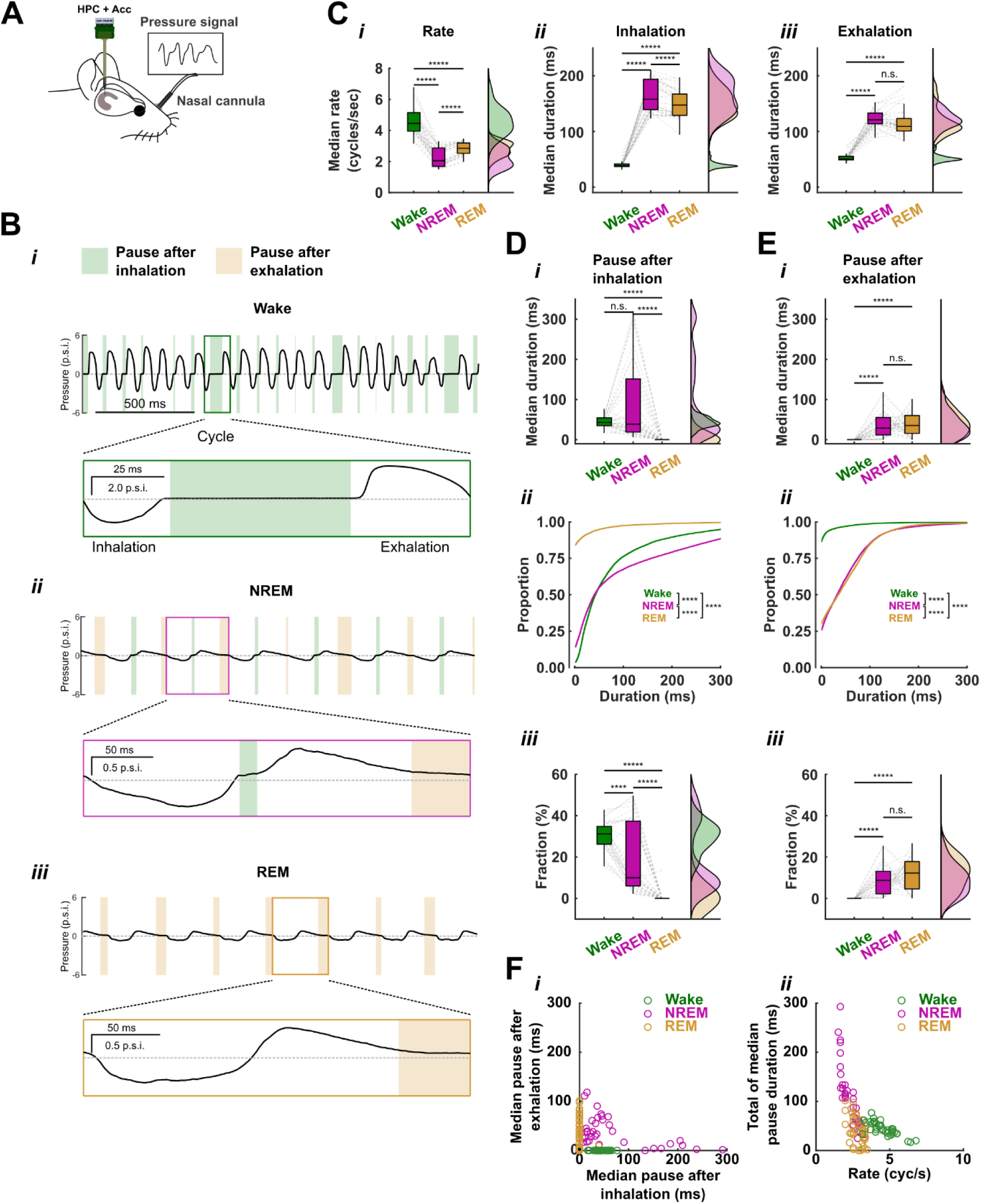
Nasal pressure recordings in freely-moving mice uncovers distinct respiratory features, including pauses, across vigilance states. **A.** Schematic representation of a mouse implanted with a nasal cannula to monitor respiration in freely-moving conditions as well as a silicon probe in the dorsal hippocampus. An accelerometer was also placed on the mouse’s headstage. **B.** Representative traces of respiration signal (2 seconds) during Wake (***i***), NREM (***ii***) and REM (***iii***) with overimposed shaded areas showing respiratory pauses after inhalation (green) and after exhalation (orange). Inhalation and exhalation correspond to negative and positive nasal pressure, respectively. Colour-coded boxes highlight one representative cycle in each states. Dashed line: 0 p.s.i. **C.** Colour-coded box plots across states (Wake = green, NREM = purple, REM = brown) showing medians of respiratory features extracted from individual recording sessions (N = 38) and their corresponding probability distributions on the right. (i) Smoothed respiration rate (bin = 200 ms, sigma = 400 ms) mean ± SEM: Wake = 4.6 ± 0.1 cycles/sec, NREM = 2.3 ± 0.1 cycles/sec, REM = 2.8 ± 0.1 cycles/sec, Kruskal-Wallis test: χ2(2) = 80.1, *P* = 4.07e-18, *post-hoc* Wilcoxon tests, *P* < 0.0001 for all. (ii) Inhalation duration mean ± SEM: Wake = 40.1 ± 1.0 ms, NREM = 164.9 ± 4.6 ms, REM = 148.6 ± 3.9 ms; Kruskal-Wallis test: χ2(2) = 78.5, *P* = 4.07e-14; *post-hoc* Wilcoxon tests: *P* < 0.0001 for all. (iii) Exhalation duration mean ± SEM: Wake = 52.4 ± 1.2 ms, NREM = 121.9 ± 2.4 ms, REM = 117.0 ± 4.0 ms; Kruskal-Wallis test: χ2(2) = 77.5, *P* = 1.50e-17; *post-hoc* Wilcoxon test: wake *vs* NREM: *P* = 7.74e-08, wake *vs* REM: *P* = 7.74e-08, NREM *vs* REM: *P* = 0.10. **D.** Duration of pauses after inhalation across states. (i) Colour-coded box plots across states showing medians of pause after inh. durations extracted from individual recording sessions (N = 38) and their corresponding probability distributions on the right. Pause after inhalation duration mean ± SEM: Wake = 44.3 ± 2.1 ms, NREM = 88.5 ± 15.7 ms, REM = 1.0 ± 1.0 ms. Kruskal-Wallis test: χ2(2) = 74.5, *P* = 6.55e-17; *post-hoc* Wilcoxon tests: Wake *vs* NREM, *P* = 0.17, Wake *vs* REM, *P* = 8.34e-08, NREM *vs* REM, *P* = 1.15e-07. (ii) Colour-coded curves representing cumulative probability distributions of the pooled pause after inhalation durations across randomly selected cycles from all 38 sessions (**Methods**). Kolmogorov-Smirnov tests with Bonferroni corrections: Wake *vs* NREM, *P* = 2.70e-118, Wake *vs* REM, *P* = 5.61e-118, NREM *vs* REM, *P* = 6.47 e-118, N = 18107 cycles for all states. Pause after inh. duration 25^th^, 50^th^, 75^th^ percentiles: Wake = 19.2, 42.4, 94.4 ms, NREM = 12.0, 40.8,157.6 ms, REM = 0.0, 0.0, 0.0 ms. (iii) Same as (***i***) for fractions occupied by pauses after inhalation relative to cycle duration. Pause after inh. fraction mean ± SEM: Wake = 30.2 ± 1.1 %, NREM = 18.3 ± 2.6 %, REM = 0.4 ± 0.4 %, Kruskal-Wallis test: χ2(2) = 80.7, *P* = 2.99e-18; *post-hoc* Wilcoxon tests: *P* < 0.0001 for all. **E.** Same as **D** for pauses after exhalation. (i) Pause after exhalation duration mean ± SEM: Wake = 0.0 ± 0.0 ms, NREM = 35.4 ± 5.0 ms, REM = 39.3 ± 4.6 ms (N = 38 sessions). Kruskal-Wallis test: χ2(2) = 68.3, *P* = 1.50e-15, N = 38; *post-hoc* Wilcoxon tests: Wake *vs* NREM, *P* = 3.64e-07, Wake *vs* REM, *P* = 1.67e-07, NREM *vs* REM, *P* = 0.76. (ii) Kolmogorov-Smirnov tests with Bonferroni corrections: Wake *vs* NREM, *P* = 1.68e-110, Wake *vs* REM, *P* = 2.39 e-110, NREM *vs* REM, *P* = 1.96e-17, N = 18107 cycles for all states. Pause after exh. duration 25th, 50th, 75th percentiles: Wake = 0.0, 0.0, 0.0 ms, NREM = 0.0, 32.8,75.2 ms, REM = 0.0, 35.2, 80.8 ms. (iii) Fraction occupied by pauses after exhalation relative to cycle duration mean ± SEM : Wake = 0.0 ± 0.0 %, NREM = 8.7 ± 1.1 %, REM = 11.8 ± 1.2 % (N = 38 sessions). Kruskal-Wallis test: χ2(2) = 70.9, *P* = 6.71e-16, *post-hoc* Wilcoxon tests: Wake *vs* NREM, *P* = 3.65e-07, Wake *vs* REM, *P* =1.68e-07, NREM *vs* REM, *P* = 0.13. **F.** Relationship between pause durations across states and their link to respiration rate. (i) Median durations of pauses after inhalation against pauses after exhalation across individual sessions (dots). (ii) Cumulative median durations of respiratory pauses (after both inhalation + exhalation) across individual sessions (dots) plotted against the median respiration rate of the same session.

To analyze the nasal pressure signals, we modified the BreathMetrics toolbox (31) so that individual respiration cycles were temporally identified and characterized in each experimental session with comparable accuracy across brain states (mouse-BreathMetrics, **Methods** and **Fig. 1B**, **Supplementary Fig. 2, 3**; **Supplementary Videos**). As expected (6, 7, 11, 14), we observed significant inter-state differences at the level of the respiration rate which was significantly more rapid during Wake (4.6 ± 0.1 cycles/sec; mean ± SEM) than during REM (2.8 ± 0.1 cycles/sec) and NREM (2.3 ± 0.1 cycles/sec), the slowest state (**Fig. 1B**, **Fig. 1C*_i_***). To exploit the precision of the pressure signal, we next focused on the dynamics of respiratory waveforms which comprised rapid alternations between negative and positive values corresponding to inhalation and exhalation, respectively (see **Fig. 1B*_i-iii_*** for representative cycles). Overall, the durations of inhalation (inh.) and exhalation (exh.) inversely scaled with the changes of respiration rates across vigilances states: inhalation was shorter during Wake compared to both NREM (*P* = 7.73e-08) and REM (*P* = 7.73e-08) and also shorter (*P* = 5.63e-07) in REM as compared to NREM (**Fig. 1C*_ii_***, inh. mean duration ± SEM: Wake = 40.1 ± 1.0 ms, NREM = 164.9 ± 4.6 ms, REM = 148.6 ± 3.9 ms) while exhalation was shorter during Wake compared to both NREM (*P* = 7.74e-8) and REM (*P* = 7.74e-8) but unchanged (*P* = 0.10) between REM and NREM (**Fig. 1C*_iii_***, exh. mean duration ± SEM: Wake = 52.4 ± 1.2 ms, NREM = 121.9 ± 2.4 ms, REM = 117.0 ± 4.0 ms).

Despite the increased peak amplitudes in Wake (**Supplementary Fig. 4Aiii**, mean inh. amplitude ± SEM: Wake = 1.81 ± 0.04 p.s.i, NREM = 0.81 ± 0.04 p.s.i., REM = 0.76 ± 0.04 p.s.i.), shorter inhalations caused overall smaller inhalation volumes (**Supplementary Fig. 4Aiv, mean inh**. volume ± SEM: Wake = 48.8 ± 1.2 p.s.i x ms, NREM = 81.6 ± 4.0 p.s.i. x ms, REM = 72.2 ± 3.5 p.s.i. x ms). In contrast, likely due to compensatory changes in exhalation amplitudes (**Supplementary Fig. 4Biii, mean exh**. amplitude ± SEM: Wake = 1.86 ± 0.04 p.s.i, NREM = 0.65 ± 0.04 p.s.i., REM = 0.78 ± 0.03 p.s.i.), changes of breathing rates across states had limited impact on exhaled volumes (**Supplementary Fig. 4Biv, mean exh**. volume ± SEM: Wake = 65.9 ± 1.1 p.s.i. x ms, NREM = 56.7 ± 2.9 p.s.i. x ms, REM = 60.6 ± 3.1 p.s.i. x ms). Overall, these data obtained with portable pressure sensors in freely moving mice confirmed the drastic change in respiratory rate between Wake and sleep and highlighted subtler differences between NREM and REM sleep.

### Respiratory pauses are a prominent feature of respiration with a signature characteristic of brain states

While respiratory waveforms exhibited well-defined inhalation and exhalation components, the high-quality intranasal pressure signals also showed flat periods corresponding to low airflow (**Supplementary Fig. 1**-3), both after inhalation (**Fig. 1B**, green shaded areas) and exhalation (**Fig. 1B**, orange shaded areas). These events were hardly detectable with intranasal thermocouples and were difficult to delineate with whole-body plethysmography (**Supplementary Fig. 1A**). Given their resemblance to respiratory “pauses” in humans (31), we referred to them as “pauses after inhalation” and ““pauses after exhalation” depending on their location within respiratory cycles. These periods were detected by mouse-Breathmetrics based on the sudden slowdown of pressure change, inducing signal inflection points used as pause onset/offset (**Supplementary Fig. 3, Methods**). Pauses occurred at both ends of inhalation and exhalation, but not inside, and were observed in all vigilance states, although examination of respiratory traces suggested differences across states (**Fig. 1B, Supplementary Fig. 2, Supplementary Videos**): during Wake, pauses predominantly followed inhalation and were practically absent after exhalation (**Fig. 1B*_i_***), whereas during REM, pauses after inhalation were absent and pauses after exhalation prevailed (**Fig. 1B*_iii_***). Pause location during NREM was in-between those two extremes with pauses still being detected after inhalation but also present after exhalation (**Fig. 1B*_ii_***).

When comparing the three states within individual sessions, the median duration of pauses after inhalation was significantly reduced during REM compared to both Wake and NREM but there was no difference between Wake and NREM (**Fig. 1D*_i_***; Kruskal-Wallis test: χ^2^(2) = 74.5, *P* = 6.55e-17, N= 38 sessions; *post-hoc* Wilcoxon signed-rank tests: Wake *vs* NREM: *P* = 0.17, Wake *vs* REM: *P =* 8.34e-08, NREM *vs* REM: *P =* 1.15e-07). However, the distributions of pauses after inhalation pooled from cycles randomly selected in all sessions (maximum of 500 cycles/session, equal number in each state, **Methods**) were significantly different across states, with NREM showing the longest pauses after inhalation followed by Wake and finally REM sleep (**Fig. 1D*_ii_***; mean duration ± SEM, Wake = 82.6 ± 0.9 ms, NREM = 105 ± 1.0 ms, REM = 9.4 ± 0.0 ms; two-sample Kolmogorov-Smirnov tests followed by Bonferroni corrections, *P <* 0.0001 for all state-pair comparisons, N = 18107 cycles for each state). This new statistical difference between Wake and NREM (as compared to session median values that were similar in **Fig. 1D*ii***) stems from a population of very long pauses in NREM (75^th^ quantile: Wake = 94.4 ms, NREM = 157.6 ms, REM = 0.0 ms). Yet, the percentage of cycle duration occupied by pauses after inhalation overall reached higher values in Wake compared to both NREM and REM (**Fig. 1D*_iii_***).

Pauses after exhalation followed an opposite pattern to pauses after inhalation when compared across states (**Fig. 1B, Supplementary Fig. 2**). As these pauses were nearly absent during Wake, their median duration extracted from individual sessions was significanty reduced compared to both NREM and REM which did not differ (**Fig. 1E*_i_***; Kruskal-Wallis test: χ^2^(2) = 68.3, *P =* 1.50e-15, N = 38 sessions, *post-hoc* Wilcoxon tests: Wake *vs* NREM: *P =* 3.64e-07, Wake *vs* REM: *P =* 1.67e-07, NREM *vs* REM: *P =* 0.76). When samples of pauses after exhalation were pooled from the 38 sessions, distributions for NREM and REM were nearly similar (**Fig. 1E*_ii_***; mean duration ± SEM, Wake = 6.0 ± 0.4 ms, NREM = 52.4 ± 0.7 ms, REM = 55.6 ± 1.0 ms; Kolmogorov-Smirnov, KS, tests followed by Bonferroni corrections, *P <* 0.0001 for all state-pair comparisons, N = 18107 cycles for each state). Similarly, the proportion of the respiratory cycle occupied by pauses after exhalation was lower in Wake compared to both NREM and REM but it was not different between NREM and REM (**Fig. 1E*_iii_***).

Together, these results suggest that the duration of respiratory pauses recapitulate distinct patterns of respiration during different vigilance states, as also shown when the median durations of pauses after inhalation and exhalation of individual recording sessions are plotted against each other (**Fig. 1F*_i_***): REM and Wake formed distinct, compact clusters close to vertical and horizontal axes, while NREM showed a large inter-session variability and spanned a large range of possible combinations. However, while the precise moment when pauses occur varies (i.e. either after inhalation or after exhalation), the fraction of the cycle occupied by pauses was ∼ 30 % for both Wake and NREM (**Supplementary Fig. 4**). In contrast, a distinctive feature of REM was the lowest fraction of the cycle occupied by pauses, only 12 %, as the proportions of both inhalation and exhalation were simultaneously extended (**Supplementary Fig. 4**).

Finally, we examined the hypothesis that long respiratory pauses are associated with low respiration rates. The relationship between the median cumulative duration of pauses within cycles and the respiration rate indeed revealed a strong negative correlation across recording sessions in each state (Wake, Pearson’s correlation coefficient, r = -0.76, *P* = 4.05e-8; NREM, r = -0.85, *P* = 1.31e-11; REM, r = -0.49; *P* = 0.002). Yet, the total duration of pauses and the respiration rate did not simply scale inversely with each other across states as measurements from all states did not fall along a single line (**Fig. 1F*_ii_***). Specifically, sleep states did show a continuum of values, which suggested scaling, but Wake stood out of the line formed by sleep sessions. This data suggested that, during sleep, episodes with longer pauses indeed correspond to slower breathing, but that other rules link respiration rate to pauses during Wake.

Our results therefore provide evidence that patterns of respiratory pauses are signatures of the three major vigilance states in freely-moving mice. These signatures did not have a straightforward link to the observed changes of respiration rates across states and therefore provide a complementary characterization of breathing-brain state relationships.

### Nasal pressure dynamics predict brain state

If respiration and brain states are intertwined, we reasoned that breathing measurements should contain sufficient information to predict whether a given cycle belongs to Wake, NREM or REM state. To test this hypothesis, we first applied a generalized linear model (GLM) to 10 respiratory features extracted from mouse-BreathMetrics (blinded for states) (**Methods**) (42 recording sessions in 7 different mice). The respiratory cycles were automatically annotated with their corresponding state (Wake, NREM or REM) thanks to the CA1 LFP signals (41) to ensure inter-session reproducibility. After normalization (log scale to attenuate large but infrequent variations, unit-standard deviation and zero-mean) (**Supplementary Table 1**), all 10 features showed a specific form of state-discrimination (e.g. inhalation and exhalation durations discriminate sleep *vs* Wake, inhalation pause duration discriminates REM *vs* other states) (**Supplementary Fig. 5A**). We therefore used them all as inputs of the prediction algorithm (**Fig. 2A**). The GLM predictions - tested on a subset of validation data excluded from the model-fitting set - were better than chance for all the three possible states, both when considering recall and precision measures (KS tests *vs* shuffled state labels for Wake: *P* = 1.89e-5, NREM: *P* = 1.89e-5 and REM: *P* = 1.89e-5 for both recall and precision, N = 10 regressions for each). For instance, REM recall was 0.84 ± 0.02, against 0.38 ± 0.02 for the shuffle (mean ± standard deviation, all values in **Supplementary Tables 2**). Yet, the three states showed different sensitivity to parameter exclusion (**Fig. 2B**) (statistics in **Supplementary Table 3**). On the one hand, Wake recall and precision were consistently high but most sensitive to the exclusion of measurements of inhalation and exhalation kinetics (Dunnett multiple test against no exclusion: *P* = 0.0000, Below Round-off Error, BRE, and *P* = 1.77e-11 for precision). On the other hand, NREM recall was most sensitive to exclusion of volume and flow features (from 0.760 ± 0.007 to 0.68 ± 0.008, Dunnett test *P* = 0.0000, BRE). Strikingly, REM recall was most impaired when pauses after inhalation were removed from the datasets (from 84% to 73%, Dunnett test *P* = 0.0000, BRE), an effect that exceeded the impact of inhalation/exhalation kinetics removal (-0.38%, Dunnett test *P* = 0.99). Removing pauses after inhalation also further deteriorated the already poor precision of REM prediction (from 23% to 16%, Dunnett test *P* = 0.0000, BRE), a reduction again more severe than when removing inhalation and exhalation kinetics (-4%, Dunnett test *P* = 1.78e-9). Thus, pauses after inhalation appeared as a key predictor of REM state, favoring recall while decreasing false positive rate, as also seen in receiving-operating characteristic curves (**Fig. 2C**).

**Figure 2.**
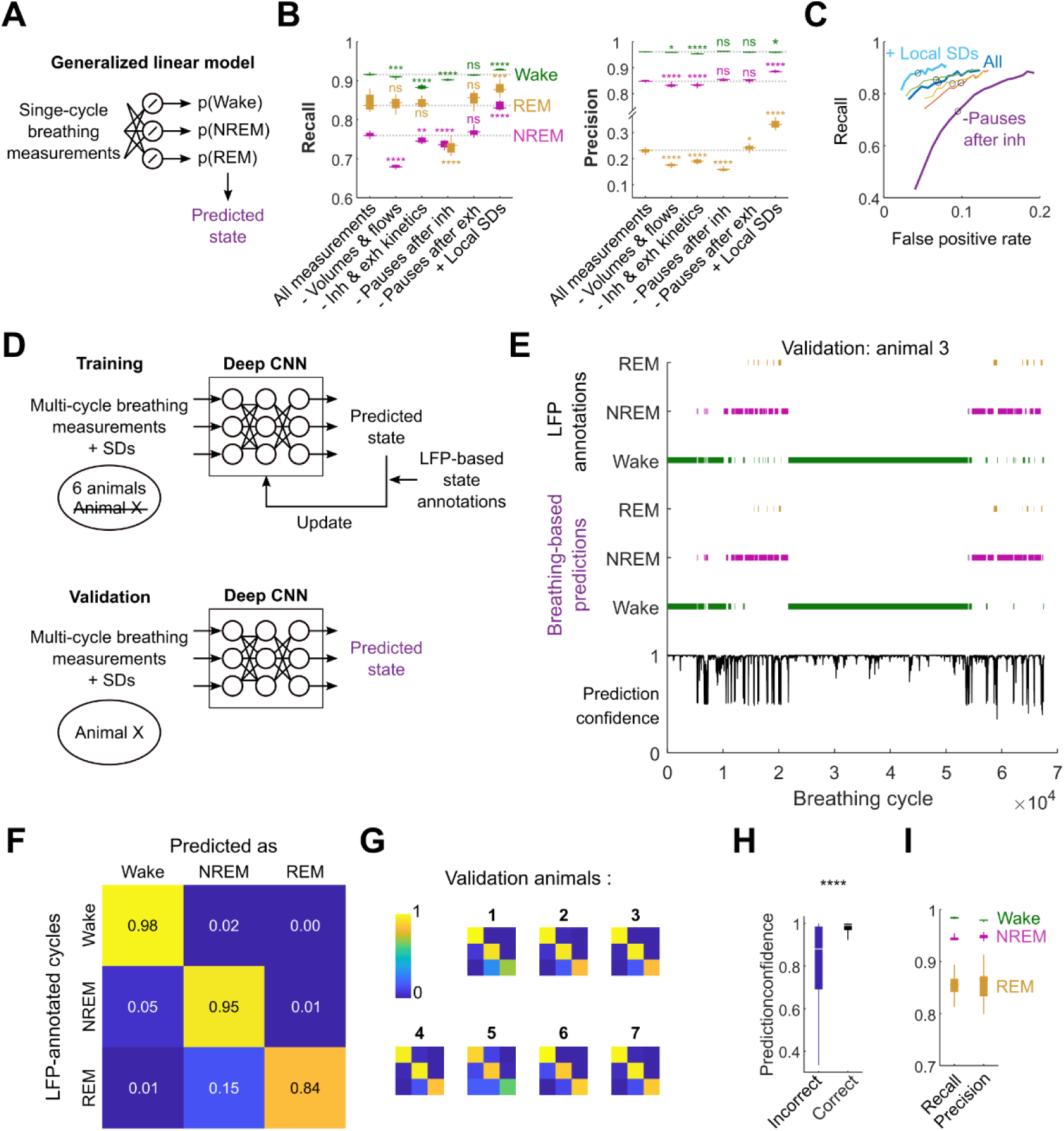
Respiration cycle properties predict brain states. **A.** Prediction of brain states based on single-cycle breathing measurements as inputs of a generalized linear model (GLM) (**Methods**). Model parameters fitting was based on the minimization of the categorical cross-entropy between the predicted states probabilities and automated annotations of brain states based on the LFP (**Methods**). **B.** Prediction performance for the model in **A** when applied to a subset of validation data not part of the model-fitting set. *Recall* = (#true positive predictions)/(#positive annotations); *Precision* = (#true positive predictions)/(#positive predictions). Either all 10 measurements were used for training and prediction (*All measurements*) or some were excluded (or added) in both training and validation steps. -*Volumes & flows*: inhalation peak flow, exhalation trough flow, inhalation and exhalation volumes were excluded. -*Inh & exh kinetics*: inhalation time to peak, exhalation time to trough, inhalation and exhalation durations were excluded. -*Pauses after inh*: exclusion of pauses after inhalation. -*Pauses after exh*: exclusion of pauses after exhalation. *+Local SDs*: additional inclusion of the standard deviations (SDs) of the 10 breathing measurements over a local time-window (10 cycles) (**Methods**). Mean ± SD values are given in **Supplementary Table 1.** Stars: Dunnett’s test *P* significance for the pairwise comparisons between exclusion groups and the ‘All measurements’ control (full statistics in **Supplementary Table 2**). N = 10 repetitions for each condition. **C.** Receiving operating characteristic of the GLM prediction algorithm for the REM state when varying the weight of REM in the objective function (lines). Black circles: balanced weights between Wake, NREM and REM states, as in **B**. Colors: different parameter exclusions conditions: dark blue = *All measurements*, red = *-Volumes &* flows, orange = *-Inh & exh kinetics*, purple = *-Pauses after inh*, green = *-Pauses after exh*, light blue = *+ Local SDs*. False positive rate = (#false positive predictions)/(#negative annotations). **D.** Supervised training strategy for the artificial neuronal network in **Supplementary Fig. 5**. Again, network parameters were adjusted to minimize the categorical cross-entropy between the predicted states probabilities and automated annotations of brain (**Methods**). For validation purpose, the training dataset included all animal recordings except for one test animal (*bottom*). All performance indicators were computed by grouping results from the 7 training paradigms associated to the 7 possible validation animals. **E.** Prediction of brain states for a session from animal #3 which was left out of the training process (validation animal). Predicted states were chosen as the ones with highest predicted probability among the three possible states (**Methods**). Prediction confidence (blue) (the maximal predicted state probability) was also provided as an index of trust in the predicted state. **F.** Median confusion matrix of single-cycle prediction of brain states when each animal (N = 7) was sequentially used for validation. Values correspond to fractions of annotations (lines sum to 1). For each validation animal, the performance indicators were computed when comparing the annotated and predicted brain states associated to all the respiratory cycles and sessions from this animal’s recordings. Those results were then averaged over 10 independently fitted networks for each validation animal. **G.** Median confusion matrices for each individual validation animal over the 10 fits. **H.** Confidence for each of the correct predictions (*Wake* as *Wake, NREM* as *NREM* and *REM* as *REM*) (n > 22e6) compared that of predictions errors (N = 723080) for the validation data used in F-G (Kolmogorov-Smirnov test *P* =0.0000, below machine precision). **I.** Median performance for the state-predicting CNN algorithm over the 7 animals sequentially left-out during training for validation (as in **D**). Boxplots characterize these median precision and recall values, for each brain state, when repeating N = 10 times the fitting procedure. Wake: recall = 0.984 ± 0.002, precision = 0.980 ± 0.002. NREM: recall = 0.943 ± 0.007, precision = 0.948 ± 0.005. REM recall = 0.853 ± 0.025, precision = 0.857 ± 0.032. Mean ± standard deviation.

So far, predictions were made on isolated cycles, ignoring their local environment and the features of the neighboring cycles. As respiration patterns are unlikely to evolve over the scale of individual cycles, we decided to complement our GLM with local standard deviations (SDs) calculated for all 10 respiration features among the 10 neighboring cycles. Local SDs showed differences across brain states (**Supplementary Fig. 5B**) but were altogether uncorrelated from the feature amplitudes (**Supplementary Fig. 5C**), which indicated that local feature variability captured another facet of the respiration-brain states relationship which could be exploited for state prediction. Indeed, by including local SDs, both prediction recall and precision were significantly improved for all the states considered (**Supplementary Tables 2 and 3**) (**Fig. 2B**). This revealed that local stability of respiration also characterizes brain states, on top of the discriminatory power of respiration kinetics for Wake, volume information for NREM, and pauses after inhalation for REM.

Even when using all breathing measurements and their local SDs, the precision for REM prediction peaked only at 0.333 ± 0.016 (mean ± SD), indicating that a majority of REM-predicted cycles are in fact mislabeled. We next aimed to decipher whether this stemmed from a limited brain state-respiration relationship, intrinsically precluding precise REM prediction, or whether this was due to the limitations of GLM approach, such as linearity and the use of single (or local) cycle measurements. To do so, we built an artificial neuronal network that is, in principle, more potent than GLM models, and trained it under supervision with the same annotated data (**Methods**). On top of the normalized respiration features, we added their SDs over a local time-window (**Methods**) as they previously improved GLM-based predictions. The resulting algorithm provides a prediction of the brain state (and a confidence indicator, as measured by the predicted state probability) associated to each single breathing cycle based on the 20 descriptors from this cycle and from its neighboring cycles (100 preceding and 100 following cycles, **Supplementary Fig. 6**). To test how efficiently our network could predict brain-states, but also how the respiration-brain state links can be generalized between animals, we trained it on 6 mice and used the left-out mouse for validation (**Fig. 2D**). As shown for an example session (**Fig. 2E**), the network learned to predict brain states for mouse #3, even though data from this animal was excluded from the training set, hence highlighting the inference of generalized, cross-animal, rules for prediction. Long periods of Wake and NREM sleep were correctly predicted with confidence close to 1 (1 corresponds to maximal confidence). Lower confidence was observed at the transition between brain states and during REM sleep periods which were much shorter, even though all 10 out of 11 REM episodes were correctly predicted.

We repeated the training procedure for all the 7 mice as “left out” (validation) animal (10 training repetitions for each animal) and assessed the prediction quality based on the median confusion matrix across animals (**Fig. 2F**). We found that 98% of annotated Wake cycles were correctly predicted as Wake cycles; 95% of annotated NREM cycles were correctly predicted as NREM and 84% of REM cycles were correctly predicted as REM. The largest confusion was observed between NREM and REM: 15% of annotated REM cycles were predicted as NREM. However, only 1% of REM cycles were incorrectly assigned to Wake and only 1% of NREM cycles were predicted as REM. Similar patterns of confusion occur when considering individual matrices associated to different validation animals (**Fig. 2G**) with minor differences which could reflect residual animal-to-animal variations that could not be compensated. Interestingly, the confidence of the predictor was the lowest when the predictions were incorrect (confidence = 0.83 ± 0.16, mean ± SD, N = 723080), while the confidence was high and showed little dispersion when annotated states matched the prediction (confidence = 0.96 ± 0.07, mean ± SD, N > 22e6) (**Fig. 2H**). Therefore, if the proposed artificial network were to be used for automated brain state annotation, prediction confidence would provide a useful indicator for downstream analysis (a feature not traditionally available for possibly faulty manual annotations and automated state prediction from LFP), encouraging caution and increased scrutiny when it is low.

We completed the analysis of prediction performance by computing standard indicators of classification performance: recall and precision (**Methods**) (**Fig. 2I**). Indicators were all ≥ 0.94 (maximum is 1) for NREM and Wake states, proving an excellent ability to recover a large fraction of the correct states (recall) but also minimal contamination of the predicted states by others (precision). REM state was still the most challenging, likely because this state’s periods are shorter and less frequent in the training sets, but indicators were still all > 0.85, indicating that the poor REM precision with GLM (**Fig. 2B, C**) stemmed from the inability of this linear, local, approach to fully exploit the rich respiration-brain state relationships. Overall, these experiments highlight the existence of general, cross-animal, rules that tightly link breathing features to brain states, and that those links are spread over multiple breathing features and their dynamics.

### Distinct co-varying respiratory features encode brain states and animal identity

The stability of performance for GLM-based state prediction obtained when sequentially excluding many, but not all, parameters (**Fig. 2B**) suggested that some features of breathing cycles co-vary as states alternate (i.e. some excluded parameters could be compensated by others), while other features behave more independently and bring distinctive information about vigilance states. In particular, pauses after inhalation seemed to uniquely discriminate REM sleep (**Fig. 2B, C, Supplementary Fig. 5A**). To characterize these co-variations, we computed the cycle-to-cycle Pearson’s correlation matrix (**Fig. 3A**) of the 10 log-normalized respiratory features characterizing respiration after balancing the proportions of brain states (1/3 of total dataset each) (**Methods**). Correlations across features showed diagonal blocks with overlaps: volume-related features (peak/trough flow and inhalation/exhalation volumes) formed one co-varying block. Kinetic measurements of inhalation and exhalation (durations, time-to-peak and - to-trough) formed another, but they showed a positive correlation with inhalation and exhalation volumes and the duration of pauses after exhalation. Interestingly, the duration of pauses after inhalation stood isolated, mainly uncorrelated with other features of respiration. Covariance matrices obtained for individual states showed similar results (**Supplementary Fig. 5C, right panels**). Principal component analysis (PCA) of the data of the combined 3 states confirmed these observations: the first principal direction (PC1), explaining more than 47.0% of the total variance (**Fig. 3B**) was mainly supported by kinetic features of inhalation and exhalation and the duration of pauses after exhalation (**Fig. 3A**). PC2 was mainly supported by volume and flow features and explained 22.9% of the total variance, and PC3 (9.64% of total variance) mainly aligned to the duration of pauses after inhalation and, to a lesser extent, to time-to-peak during exhalation.

**Figure 3.**
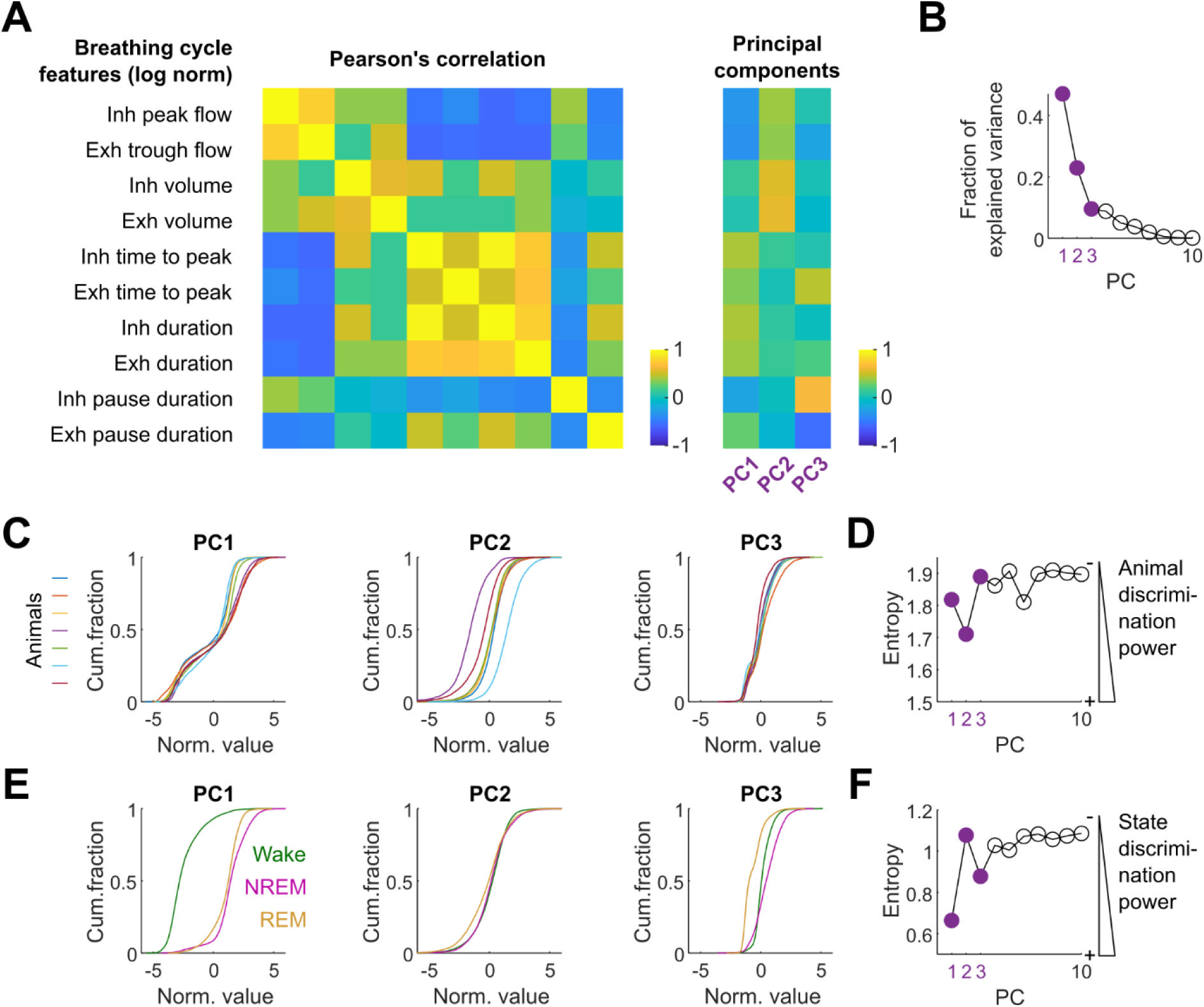
Distinct respiration features co-vary with brain states and animal identity. **A.** Correlation between features of single respiration cycles (N = 42000) after log-normalization (**Supplementary Table 1**) and the first three principal components (PCs) obtained by PCA. Proportion of brain states (Wake/NREM/REM) and animal identities (7 mice) were equalized. Inh: inhalation. Exh: exhalation. **B.** Fraction of the total data variance explained by all PCs. The first three largest are denoted PC1, 2 and 3 in decreasing order of explained variance. **C.** Cumulative distributions of coefficients associated to the PC1 2 and 3 (A) for the individual 7 mice (N = 6000 for each). **D.** Entropy of single cycle PC coefficients computed from *Pr(animal id|coefficient value)* (**Methods**) for all PCs. Lower entropy indicates better identity predictive power. **E.** Cumulative distributions of coefficients associated to Wake (N = 14000), REM (N = 14000) and NREM (N = 14000) states for all animals pooled together. **F.** Entropy of single cycle PC coefficients computed from *Pr(brain state|coefficient value)* (**Methods**) for all PCs. Lower entropy indicates better identity predictive power.

We next investigated two factors which could underlie the three first PCs by contributing to data diversity: brain state differences, but also animal-to-animal variations. Two-way analysis of variance (ANOVA) of PC coefficients revealed that those two factors and their interaction all significantly contributed to data diversity for the three first PCs (for all PCs *P* = 0.000, BRE, for the state and animal identity factors and their interactions). However, animal identity and brain states had different weights: for both PC1 and PC3 coefficients the *F*-statistics was much larger for the brain state factor (*F*_state_ = 41896, degrees-of-freedom df = 2; *F*_animal_ = 244, df = 6; *F*_state x animal_ = 207, df = 12 for PC1; *F*_state_ = 8552, df = 2; *F*_animal_ = 309, df = 6; *F*_state x animal_ = 248, df = 12 for PC3) showing its dominance, while animal identity was the most prominent factor for PC2 coefficients (*F*_state_ = 335, *F*_animal_ = 3690, *F*_state x animal_ = 138). In good agreement with the ANOVA, the individual distributions of coefficients for the 7 analyzed animals overlapped prominently for PC1 and PC3, but clear animal-to-animal differences were visible for PC2 (**Fig. 3C**). This was confirmed when computing the predictive power of individual PC coefficients with respect to animal identity: PC2 entropy (**Methods**) was 5.1% and 9.5% lower (lower is more predictive) than that of PC1 and PC3, respectively (**Fig. 3D**). Unlike animal identity, brain states were clearly discriminated by PC1 and PC3, but not PC2, when considering the distribution of individual coefficients (**Fig. 3E**) or the state-predictive power as measured by entropy (+62.3% entropy increase from PC1 to PC2) (**Fig. 3F**). The most striking difference for PC1 coefficients was between Wake *vs* sleep states, unlike PC3 which mainly differentiated REM vs NREM+Wake (**Fig. 3E**), hence highlighting differences in brain state coding for those feature components. The entropies for other, less representative PCs (**Supplementary Fig. 7A**) were not as marked as for PC1, PC2 and PC3 (**Fig. 3D, F**). Overall the analyses of single cycle covariates revealed that: 1) kinetic features of inhalation and exhalation contributed the most to the variability of the data (best represented by PC1), were robust across animals and discriminated sleep and Wake states, 2) flow and volume features (best represented by PC2) varied across animals but less across states and 3) changes in pauses after inhalation (best represented by PC3) were an distinctive feature of REM sleep, consistent with the degraded GLM-based prediction of REM when pauses after inhalation are excluded (**Fig. 2C**).

### Progressive elimination of pauses after inhalation anticipates transition to REM sleep

Using PC1 and PC3 to summarize breathing features, we plotted time-series of coefficient values against that of brain states for an exemplar session (**Fig. 4A**). In agreement with the population analysis (**Fig. 3**), PC1 and 3 coefficients reached different plateaus when the cortical state changed and PC3 showed larger differences between the plateaus surrounding the transitions to/from REM. Yet, instead of instantaneous changes or respiration synchronized with state transitions, examples #1-6 hinted that coefficient adaptation dynamics might differ between types of transitions and between PC1 and 3. We therefore asked next how tightly breathing is linked to brain states through time by characterizing breathing adaptation in narrow time windows around state transitions.

**Figure 4.**
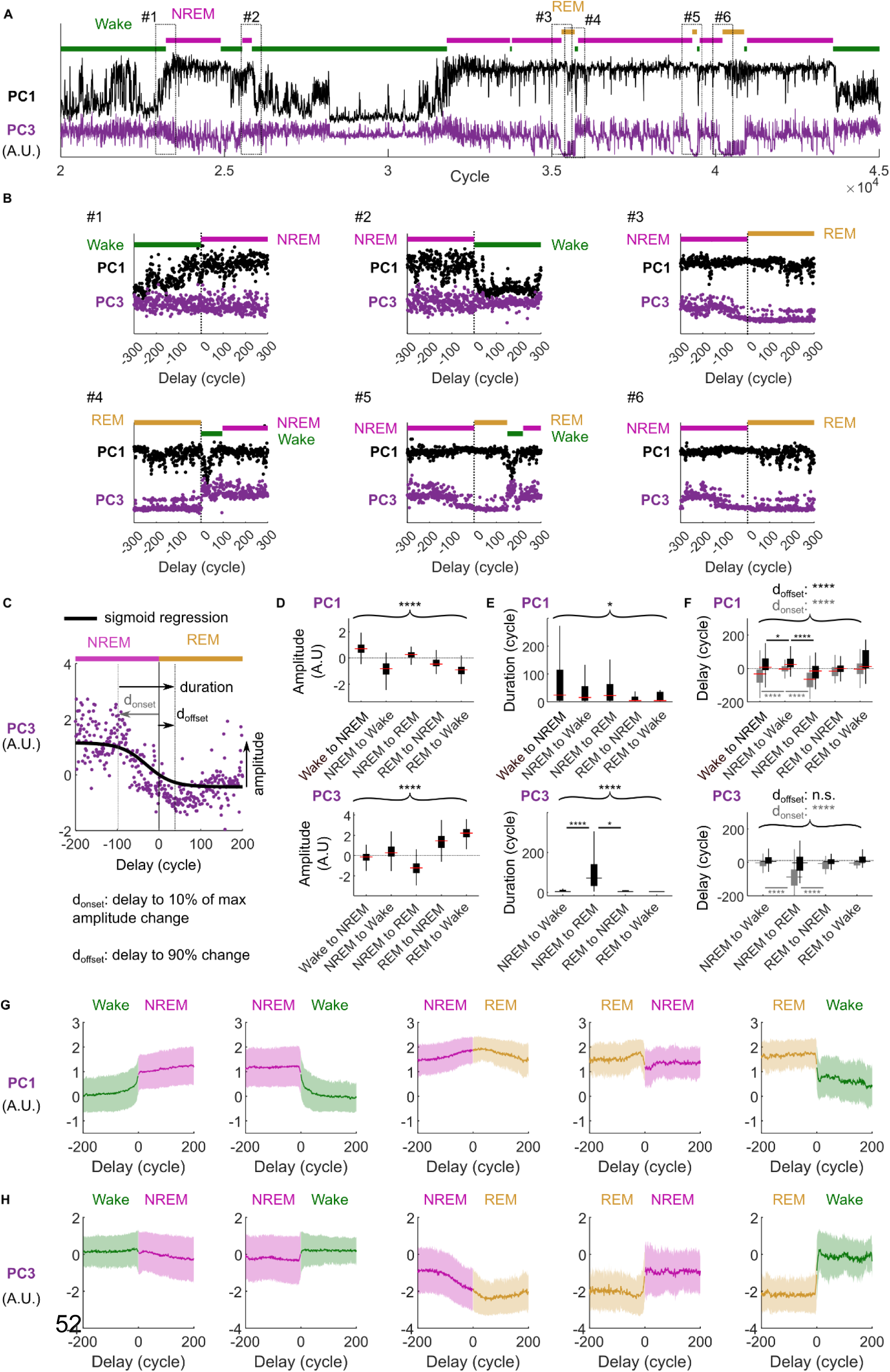
Fast and slow anticipatory adaptation of breathing at Wake→NREM and NREM→REM transitions. **A.** Time series of PC1 and 3 coefficients for single-cycle measurements aligned to annotated brain states for an exemplar session. **B.** Zoom in on size transition events indicated in A. **C.** Sigmoid function regression of PC coefficients around an example transition from NREM to REM sleep. Based on the sigmoid parameters, the amplitude of the PC value change, the onset of the adaptation (d_onset_ = delay between the time point of 10% of amplitude max change and the time of the annotated state transition) and its offset (d_offset_ = delay for 90% amplitude change) were computed. The duration of adaptation was computed as duration = d_offset_ - d_onset._ For the sake of robustness, only transitions which showed a statistically significant change of values were fitted with a sigmoid function (**Methods**). For non-significant changes, the amplitude was computed as the difference between the PC values post (300 cycles) and pre (300 cycles) state transitions. Number of measurements computed in each case are summarized in **Supplementary Table 4**. Mean ± SD of transition parameters for all the transition types are given in **Supplementary Tables 5** and **6**. **D.** Distribution of the amplitude changes for PC1 and 3 at state transitions. ANOVA statistics in **Supplementary Table 7**. Statistics of pairwise comparisons between all state transitions are given in **Supplementary Table 8**. **E.** Comparison of the adaptation duration parameter at state transitions. For PC3, the low change of coefficients amplitude at Wake→NREM transitions precluded robust sigmoid fitting. Results from ANOVA with the other transitions are in **Supplementary Table 7**. Statistics of pairwise comparisons between all state transitions are in **Supplementary Table 9**. **F.** Same comparisons as in **E**, but for the adaptation onset (d_onset_) and offset (d_offset_) parameters. Results from ANOVA with the other transitions are in **Supplementary Table 7**. Statistics of pairwise comparisons between all state transitions are in **Supplementary Tables 10**. **G.** Population averages of PC1 coefficients for all state transitions in the recording sessions from 42 sessions of 7 animals. Wake→NREM N = 672; NREM→Wake N = 524; NREM→REM N = 232; REM→NREM N = 92, REM→Wake N = 141. Plot: average value (solid line) ± standard value (shade). Cycles that did not match the studied states because of extra state transitions in the analysis window were clipped. **H.** Same as **G**, but for PC3 coefficients.

In example #1 (**Fig. 4B**), PC1 adaptation preceded Wake→NREM transition by more than 50 respiratory cycles, a feature also observed in examples #4 and #5. In a mirror fashion, PC1 adaptation seemed to follow transition back to Wake in example #2. To precisely quantify these dynamics, we computed the coefficient amplitude change at each individual state-transition in all recording sessions and fitted a sigmoid function to coefficient values when the change was statistically significant (**Methods**) (**Fig. 4C, D**). It allowed us to derive 3 characteristic parameters: the time delay of adaptation onset as compared to the state change time point (d_onset_), its offset (d_offset_), and the duration of adaptation (d_offset -_ d_onset_) (**Fig. 4E, F**, top row) (N and mean ± SD for all parameters in **Supplementary Tables 4-6**). One-way ANOVA confirmed that, indeed, different transitions showed discriminating dynamics of respiratory adaptation for PC1 (**Supplementary Table 7**) (**Fig. 4D-F**, top row). In agreement with examples in **Fig. 4B**, the largest changes of amplitudes were seen at sleep/wake transitions (e.g. Wake→NREM: 0.75 ± 0.51 A.U., NREM→Wake: -0.88 ± 0.59, NREM→REM: 0.24 ± 0.33, mean ± SD A.U., in **Supplementary Table 8** all pairwise tests *P* < 1e-4, except for NREM→Wake vs REM→Wake). This large change in PC1 amplitude at transition to/from sleep was also clearly visible when averaging coefficients from all transitions in all the 42 recording sessions (**Fig. 4G**). Similarly to examples #1, 2, 4 and 5, systematic sigmoid fitting revealed that PC1 coefficients started to adapt before the annotated Wake→NREM transition and stabilized around it: d_onset_ = -54.5 ± 69.6 cycles, d_offset_ = 19.8 ± 78.1 cycles (N = 343; mean ± SD), while the adaptation onset synchronized with the transition back to Wake: d_onset_ = -6.2 ± 49.1 cycles, d_offset_ = 42.6 ± 59.6 cycles (N = 340; mean ± SD) (Wake→NREM *vs* NREM→Wake Tukey’s honestly significant difference, HSD, *P* = 0.0000, BRE for d_onset_ and *P* = 0.0167 for d_offset_, **Supplementary Table 10**). Respiratory transition to sleep was also 52% more progressive than from sleep (Wake→NREM duration = 74.3 ± 97.1 cycles, NREM→Wake 48.8 ± 73.5 cycles, Tukey’s HSD *P* = 0.03). Notably, dynamics of individual respiratory features associated to PC1 (inhalation and exhalation kinetics) matched these dynamics at state transitions (but not as saliently) (**Supplementary Fig. 8 E-F**). PCs that poorly discriminated brain states (including PC2) generally did not show clear adaptation at state transitions (**Supplementary Fig. 7B**).

In contrast with PC1, PC3 coefficients were rather stable at Wake↔NREM transitions in examples #1, 2, 4 and 5 (**Fig. 4B**), at the population level (**Fig. 4H**), and when computing individual amplitude coefficients (Wake→NREM: -0.22 ± 0.51, N = 672; NREM→Wake: 0.44 ± 0.74, N = 524) (**Fig. 4D**). Yet, the amplitude of the adaptation depended on the type of state transition (one-way ANOVA, *P* < 1e-4, **Supplementary Table 7**). Pairwise comparisons revealed that the largest increases of PC3 values occurred when exiting REM sleep (to NREM: 1.34 ± 0.90, to Wake: 2.13 ± 0.76), while the largest decrease occurred when entering it (-1.21 ± 0.77) (**Supplementary Table 8**). These patterns of PC3 coefficients matched the lack of adaptation of pauses after inhalation (as these contribute most prominently to PC3, **Fig. 3A**) at Wake↔NREM transitions and their elimination when entering REM (**Supplementary Fig. 8I**). Strikingly, respiratory adaptation to REM frequently outlasted 100 cycles (NREM→NREM duration = 91.26 ± 79.45 cycles, N = 192) and mainly occurred in the time window preceding the annotated transition: d_onset_ = -96.41 ± 85.10 cycles, d_offset_ = -5.14 ± 63.64 cycles. This was clearly seen in examples #3 and 6 in **Fig. 4B** as PC3 slowly decreased before transition to REM, and at the population level with a diagonal curve of PC3 coefficients starting ∼100 cycles before transition to REM and plateauing shortly into it (**Fig. 4H**). The reverse changes when exiting out of REM were more abrupt (REM→NREM duration = 51.20 ± 103.90 cycles, N = 92; REM→Wake duration = 46.40 ± 104.74 cycles, N = 141; Tukey’s HSD *vs* NREM→REM *P* = 0.04 and 0.09 respectively) (examples #4 and 5 in **Fig. 4B**, population averages in **Fig. 4H**).

Overall, our data indicate that distinct respiratory changes presage the transition to deeper sleep states. Increasing PC1 coefficients, corresponding to slower breathing, shortly (∼15 s for a 3 Hz breathing rate) preceded Wake→NREM transitions. More surprisingly, a progressive reduction in the duration of pauses after inhalation (reflected by PC3) consistently preceded REM entries by about 30 s. Reverse transitions back to Wake or NREM were associated with the reverse respiratory changes, but they followed the annotated state transitions at an accelerated rate, as if the transitions were unanticipated.

### Mirrored ramping of respiratory pauses during NREM packets

So far, our data showed that distinct features of respiration tightly track changes in cortical states at transitions between Wake, NREM and REM. Given this relationship, can respiration track finer state transitions such as the micro-arousals (MAs) that delineate the spindle-rich packets tiling NREM sleep (20–22)? To address this question, we identified MAs as before (42) as transient episodes of high acceleration variation (**Methods**) in all recording sessions (n = 3022 MAs, mean ± SEM duration = 9.8 ± 0.3 seconds) and used them to delineate NREM packets (**Methods**) (**Fig. 5A_i_**). NREM packets preceding/following Wake and REM were excluded as transitions with Wake or REM were previously characterized (**Fig. 4**). There were a total of N = 2242 NREM packets in our dataset (frequency = 0.83 per minute) and the durations and spectral properties of these packets matched with previous reports (20–22): the durations spread over two orders of magnitude with a median of 38 seconds (**Fig. 5B**) and the power in both sigma (10-20 Hz) and delta (0.5-4 Hz) frequency bands of LFP fell off at their edges (**Fig. 5C**). This was evident in an example long stretch of NREM sleep (**Fig. 5A**) where MAs were temporally aligned to power troughs for delta and sigma bands (**Fig. 5A_ii_**). Strikingly, MAs delineating packets also seemed to be narrowly aligned to transient increases and decreases of the durations of pauses after inhalation and exhalation, respectively (**Fig. 5A_i_**). This was particularly apparent when analyzing data pooled from all packets: in normalized time pauses changed abruptly the edges of packets (**Fig. 5C_ii_**) and the duration of pauses after inhalation abruptly increased ∼5 s before packet offset (i.e. MA onset) (**Fig. 5D_ii_**) while pauses after exhalation dropped (**Fig. 5D_iii_**). We also found that other respiratory features changed at MAs (**Supplementary Fig. 10B**), indicating that MAs during NREM are tightly associated to a general change of respiration mode that precedes movement twitches.

**Figure 5.**
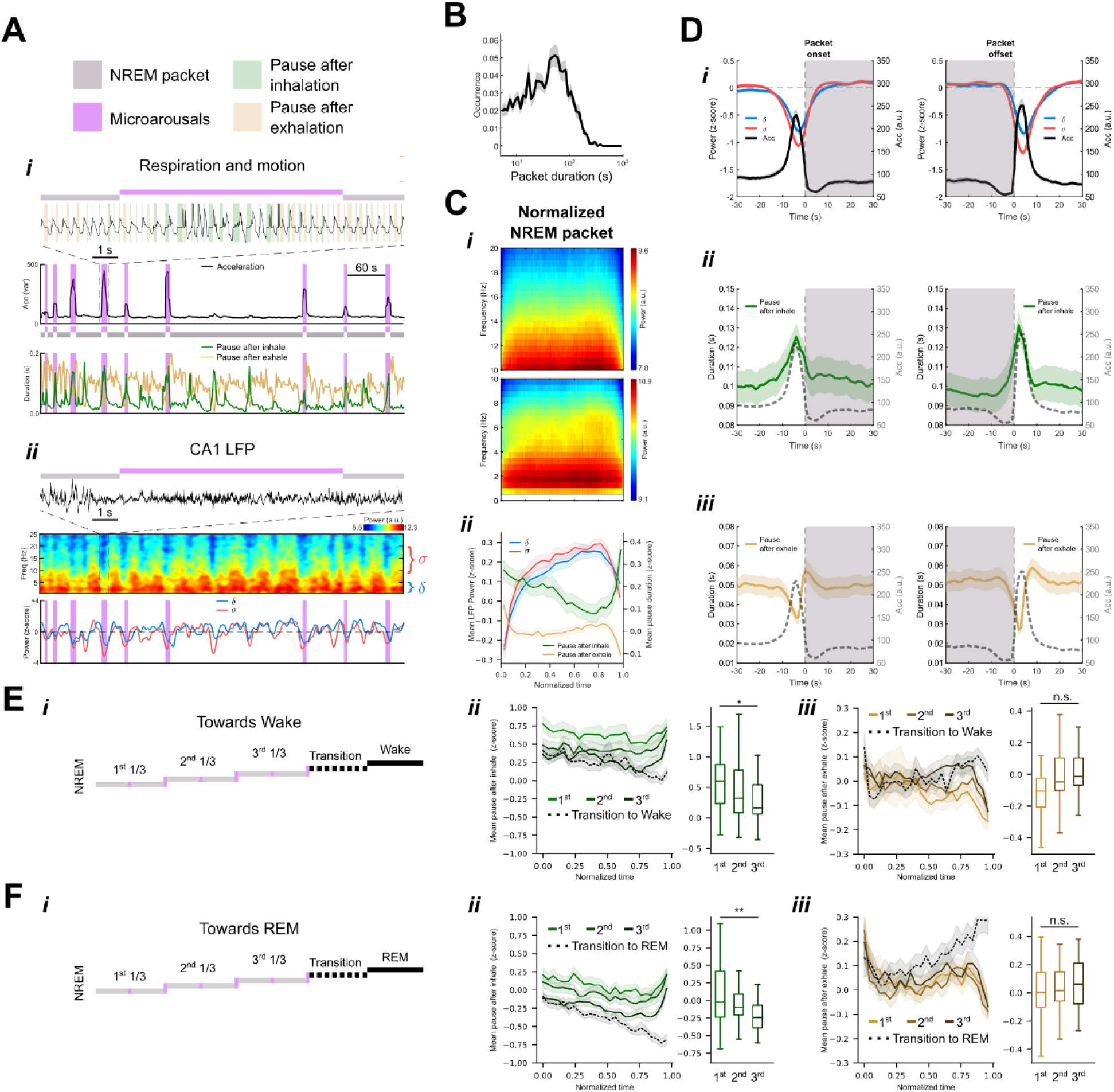
Sharp changes in respiratory pauses signal microarousals/NREM packet transitions and flucturations in hippocampal LFP. **A.** Coordination between respiration, motion and hippocampal LFP during representative example of NREM sleep. ***i***) Typical structure of NREM sleep showing the alternation of long epochs of immobility (NREM packets, light gray), and short transient movements (micro-arousals, dark purple) as revealed by the local standard deviation of acceleration signal. ***ii***) Concomitant hippocampal spectrogram during ***i*** and relative delta/sigma power fluctuation. **B.** Mean (line) + SEM (shaded area) histogram of NREM packet duration (*x*-axis in log_10_ scale) shows consistent peak between 10-100 seconds. **C.** (***i***) Averaged spectrogram across sessions (N = 38) during normalized NREM packets (50 non-overlapping time-bins) for sigma (top) and delta (bottom) frequency bands. (***ii***) Mean (line) + SEM (shaded area) of z-score delta/sigma power and pause duration. 25 non-overlapping time-bins were used for time normalization across time windows. **D.** Coordination between hippocampal LFP correlates, motion and respiratory pause patterns during NREM packet (light gray area) at transitions (onset = left column, offset = right column). ***i***) Mean (line) + SEM (shaded area) z-scored delta/sigma power dropping during acceleration peaks (i.e. micro-arousals). Mean (line) + SEM (shaded area) z-scored durations of pause after inhalation (***ii***) and pause after exhalation (***iii***). Dashed grey line: acceleration signal. **E.** (***i***) Grouping in 3 terciles of packets from an NREM epoch leading to Wake according to their chronological order. A last time window, ‘Transition’, corresponded to the time-period between the last MA and the transition to subsequent Wake. (***ii***) and *(**iii***) average duration of pauses after inhalation and exhalation (z-scores computed over NREM period) across sessions N = 36 (mean ± SEM). 25 non-overlapping time-bins were used for time normalization across time windows. Boxplots: comparison of the z-scores averaged over the 25 time-bins across sessions. One-way ANOVA between terciles: *P* = 1.56e-2 for pauses after inhalation, *P* = 0.47 for pauses after exhalation. Post-hoc comparisons between terciles for pauses after inhalation: 1^st^ vs 2^nd^ *P* = 0.2311, 1^st^ vs 2^nd^ *P* = 0.0113, 2^nd^ vs 3^rd^ *P* = 0.40. N = 36 sessions for each group. Only epochs with at least 3 NREM packets were included, hence discarding 2 sessions without such an epoch. **F.** Same analyses as for **E**, but for NREM epochs that are immediately followed by an REM episode. One-way ANOVA between terciles: *P* = 6.23e-3 for pauses after inhalation, *P* = 0.83 for pauses after exhalation. Post-hoc comparisons between terciles for pauses after inhalation: 1^st^ vs 2^nd^ *P* = 0.3600, 1^st^ vs 2^nd^ *P* = 1.9e-3, 2^nd^ vs 3^rd^ *P* = 9.62e-2. Only epochs with at least 3 NREM packets were included, hence discarding 4 sessions without such an epoch.

Besides MA, we found that within packets, the average duration of pauses after inhalation continuously decreased while that of pauses after exhalation showed an inversed dynamic in the second half of the packets (**Fig. 5C_ii_**). Other respiratory features were more stable (**Supplementary Fig. 10A**). Interestingly, when comparing packets, we found that the durations of pauses after inhalation (and not pauses after exhalation) was not homogeneous across NREM epochs and packets (**Fig. 5E, F**). On average, packets in epochs that were followed by a REM episode had lower duration of pauses after inhalation (mean z-score = -0.09 ± 0.52, mean ± SD, N = 1048 packets) than those, presumably of lighter sleep, that led back to Wake directly (mean z-score = 0.41 ± 0.81, mean ± SD, N = 1118 packets) (KS test *P* = 2.14e-38) (**Fig. 5E*_ii_*** vs **Fig. 5F*_ii_***). This suggested that the prominence of pauses after inhalation is linked to the depth of sleep. Indeed, when we grouped packets in three categories according to their chronological order of occurrence in NREM epochs (1^st^, 2^nd^ and 3^rd^ terciles), there was a decline of the mean duration of pauses after inhalation across the three terciles whether the epoch was followed by Wake (**Fig. 5E*_ii_***) (one-way ANOVA *P* = 1.56e-2) or REM (one-way ANOVA *P* = 6.23e-3) (**Fig. 5F*_ii_***). This was not the case for the duration of pauses after exhalation which remained stable throughout NREM packets (**Fig. 5E*_iii_***, **Fig. 5F*_iii_***). Moreover, the lowest level of pauses after inhalation occurred in the 3^rd^ tercile before REM, presumably the deepest stage of NREM sleep. The dynamics of pauses there recalled that of the ensuing period of transition to REM, except for the rapid increase of pause duration at the end of the packet (just before MA) which was not seen when the circuit transitioned to REM: without MA, the duration of pauses after inhalation continued to decrease further (**Fig. 5*_ii_***). These results suggest that pauses after inhalation reflects the progression to deeper sleep stages and could explain median variability observed across sessions (**Fig. 1**) as sleep patterns differed in between animals and recording days.

### Sigma power troughs during NREM sleep are associated with abrupt changes of respiratory pauses

If a sudden drop of sigma-band LFP-power coincided with a sharp increase of pauses after inhalation during MAs in NREM sleep (**Fig. 5C_ii_, D_ii_**), the examination of exemplar recording sessions revealed that this inverse relationship may also hold true outside of movement-defined MAs (**Fig. 5A**, **Fig. 6A**). In fact, when we identified troughs in sigma-band power (**Methods**), they temporally aligned with pauses after inhalation even when no animal motion was detected (**Fig. 5A**, **Fig. 6A**). To quantify this phenomenon, we examined the evolution of pauses after inhalation and after exhalation around sigma troughs concomitant with (MA^+^) or without (MA^−^) micro-arousals (**Fig. 6B-D**). This revealed that a few seconds before sigma (and delta, **Fig. 6Biii**) power reached a trough, pauses after exhalation shortened whether a MA occurred or not (MA^+^ = 31 ± 3 ms, MA^−^ = 37 ± 5 ms, mean across sessions ± SEM, N = 38 sessions). These pauses were barely shorter when sigma troughs co-occurred with MA (MA^+^, Wilcoxon signed-rank test, *P* = 0.02, N = 38) (**Fig. 6C**). For MA^+^, this drop in pause after exhalation duration was followed by a sudden elongation of these pauses in the 5-10 seconds following the trough (**Fig. 6C**) in agreement with results for NREM packet offsets (**Fig. 5C, D**). Elongation of these pauses also occurred after sigma troughs without MA yet, with a slower dynamic and no overshoot (MA^+^ vs MA^−^ power at 5s post trough (MA^+^ = 57 ± 6 ms, MA^−^ = 48 ± 5 ms, mean across sessions ± SEM, N = 38 sessions, Wilcoxon test, *P* = 0.003). These results suggest that shortening of pauses after exhalation are tied to sigma power troughs, while the following overshooting rebound of pause duration is instead linked to the physiology of MAs.

**Figure 6.**
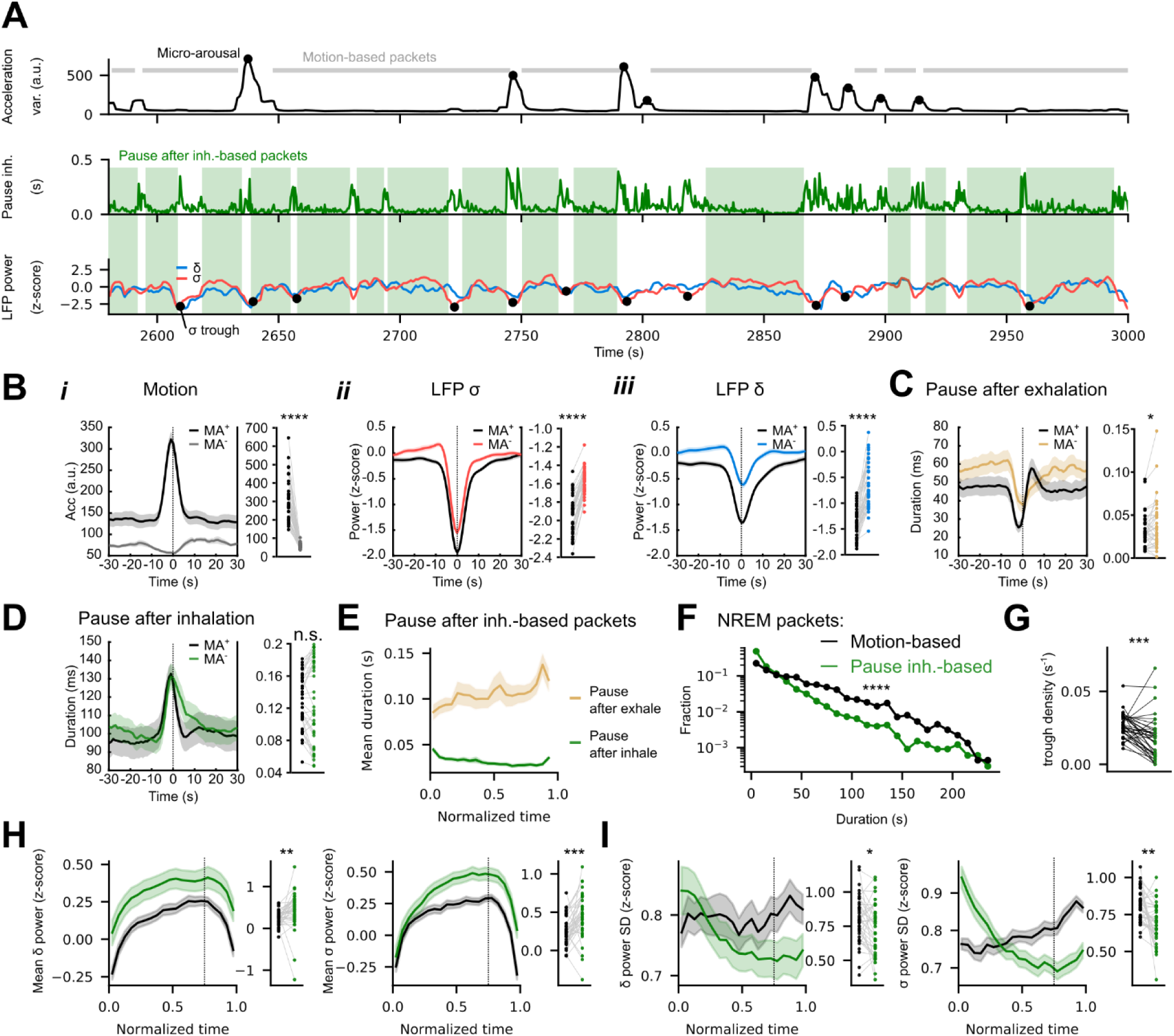
Bouts of pauses after inhalation align with sigma-band power troughs and delineate respiration-based packets in NREM. **A.** Example of 7 min of NREM sleep with concurrent recording of the animal motion (top), pauses after inhalation (middle) and LFP power in delta and sigma bands (bottom). Micro-arousals (top panel, black dots) were defined as transient periods of large motion, while large drops in the sigma-band power defined ‘σ troughs’ (bottom panel, black dots) (**Methods**). NREM packets have been defined either as periods of low motion (grey bars) delineated by micro-arousals, or, for the first time, as periods devoid of pauses after inhalation (green boxes) (**Methods**). **B.** (*Left*) Mean (line) + SEM (shaded area) peri-event curves (N = 38) centered on sigma troughs of acceleration (***i***), LFP power (z-score during NREM) for sigma-(***ii***) and delta-band (***iii***), concomitant with (MS^+^, black) or without micro-arousals (MS^−^, color-coded). Right panels: comparisons between MS^+^ and MS^−^ at t = 0 (sigma troughs time point) (Wilcoxon signed-rank tests across sessions). Acceleration signal (a.u): MA^+^ = 314 ± 18, MA^−^ = 60 ± 2, *P* = 7.74e-8. Sigma power (z-score): MA^+^ = -1.95 ± 0.03, MA^−^ = 1.58 ± 0.02, *P* = 7.74e-8. Delta power (z-score): MA^+^ = -1.36 ± 0.05, MA^−^ = -0.66 ± 0.07, Wilcoxon test across sessions, *P* = 1.25e-7. **C.** Same as B for the duration of pauses after inhalation. **D.** Same as B for the duration of pauses after exhalation. **E.** Mean ± SEM duration of pauses after inhalation and exhalation in normalized time (20 time bins) over NREM packets defined by epochs with low pauses after inhalation (**Methods**). Values averaged first per session. N = 38 sessions. **F.** Histograms (5s bins) of the duration of motion- and pauses after inhalation-based NREM packets. Fractions in log-scale to highlight the differences in the long-tail of the distributions. Motion-based packets: mean = 49.93 s, SD = 43.96 s, 25^th^ quantile = 16.50 s, median = 38.17 s, 75^th^ quantile = 70.31 s, N = 2242 packets. Respiration-based packets: mean = 27.19 s, SD = 35.43 s, 25^th^ quantile = 8.0 s, median = 15.5 s, 75^th^ quantile = 31.5 s, N = 3488 packets. Wilcoxon-Mann-Whitney test *P* = 2.43e-143. **G.** Mean density of sigma troughs per packet in the same recording sessions. Wilcoxon test across sessions: statistics = 118, *P* = 2.50e-4, N = 38 sessions. Motion-based packets: mean ± SD = 0.026 ± 0.008 troughs/s. Respiration-based packets: mean ± SD = 0.018 ± 0.014 troughs/s. **H.** Mean ± SEM of the mean (left) of the delta- and sigma-band power across sessions in normalized time (20 bins) trough NREM packets. Black: motion-based packets. Green: pause after inhalation-based packets. N = 38 sessions. Boxplots: respiration- and motion-based packet values for the 75^%^ normalized time bin (dashed line). At the 75^%^ normalized time bin: motion-based packets = 0.25 ± 0.18 for delta, 0.29 ± 0.16 for sigma, respiration-based packets: 0.41 ± 0.44 for delta, 0.48 ± 0.31 for sigma (mean ± SD). Wilcoxon test across sessions delta power: *P* = 1.87e-3, sigma power: *P* = 2.97e-4; N = 38. Z-score normalization was computed over NREM cycles. **I.** Same as G but for the standard deviation (SD) of the LFP power z-scores in individual sessions. At the 75^%^ normalized time bin: motion-based packets = 0.80 ± 0.17 for delta, 0.84 ± 0.11 for sigma, respiration-based packets: 0.72 ± 0.19 for delta, 0.69 ± 0.14 for sigma (mean ± SD). Wilcoxon test across sessions: *P* = 1.71e-2 for delta, *P* = 2.11e-3 for sigma; N = 38. Z-score normalization was computed over NREM cycles.

In the opposite direction, pauses after inhalation elongated in a similar fashion whether MA occurred or not and they had similar durations at sigma troughs in presence or absence of MA (MA^+^ = 128 ± 6 ms, MA^−^ = 128 ± 8 ms; Wilcoxon test, *P* = 0.77, N = 38 sessions) (**Fig. 6D**). Similar to the other category of pauses, the dynamic observed after the trough was slower in absence of MA (MA^+^ vs MA^−^ power at 5s post trough: MA^+^ = 98 ± 8 ms, MA^−^ = 109 ± 9 ms; Wilcoxon test, *P* = 0.77, N = 38 sessions, Wilcoxon test *P* 0.0003, N = 38), as was the absence of overshoot in this condition. These observations suggest that the link between pauses and sigma-band power is general to the NREM state and goes beyond the specific physiology of MAs.

### Long pauses after inhalation delineate homogeneous packets of high sigma power

Some periods of NREM sleep showed clear segments during which pauses after inhalation were absent and were delineated by bouts of pauses that coordinated with sigma-band troughs (**Fig. 6A**). We therefore identified novel NREM packets (‘respiration-based packets’) defined by the absence of pauses after inhalation (**Fig. 6A**, middle-row, green boxes) - rather than on animal motion - by simply applying a threshold (75 ms) to the duration of pauses after inhalation (**Methods**). By doing so, we obtained NREM packets with homogeneously low pauses after inhalation and correspondingly, high pauses after exhalation (**Fig. 6E**). Those packets were shorter (27.2 *vs* 49.9 s mean duration) and more homogeneous in duration (35.4 *vs* 44.0 s SD of durations) than the more-traditional ones based on motion (Wilcoxon test between durations *P* = 2.43e-143, respiration-based packets N = 3488, motion-based packets N = 2242). In particular, very long packets (>1 min) of low motion were replaced by shorter packets with no (or small) pauses (**Fig. 6F**). This is exemplified in **Fig. 6A** where a standard packet of ∼100 s was divided in 5 shorter respiration-based packets whose ends were often associated to sigma power troughs (it was not the case for two packets but sub-threshold sigma power inflections were observed at the ends of these packets nonetheless). Hence, and as expected from our findings in **Fig. 6D, E**, the newly defined packets contained fewer sigma troughs per second (-31% mean density per session, Wilcoxon test: *P* = 2.50e-4, N = 38) (**Fig. 6G**). This suggests that respiration-based packets delineated more homogeneous periods of LFP power. For confirmation, we temporally aligned pause after inhalation-based packets and plotted the mean delta and sigma-band LFP power in normalized time (**Fig. 6H**). For both bands, power z-scores increased at the start of the packet, reached a maximum value at ∼75% of the packet length, and then dipped towards lower values at the end. Importantly, averaged z-scores at the 75%-time point of the packets were greater than for motion-based packets for both sigma (power z-score = 0.48 ± 0.31 for respiration-based packets, 0.29 ± 0.16 others, mean ± SD, Wilcoxon test *P* = 2.97e-4, N = 38) and delta (0.41 ± 0.44 *vs* 0.25 ± 0.18, Wilcoxon test *P* = 1.87e-3, N = 38) bands (**Fig. 6H**). When considering the variability of delta and sigma z-score in packets, the mean standard deviation was significantly reduced for sigma at the 75%-time point as compared to motion-based packets (-18%, Wilcoxon test *P* = 2.11e-4, N = 38), while SD of delta showed a more modest decrease (-10%, Wilcoxon test *P* = 1.87e-3, N = 38) (**Fig. 6I**). This again confirmed that the transient drops in sigma power are confined to the edges of respiration-based packets (where pauses after inhalation occur), while sigma power is less stable and generally lower throughout motion-based packets.

### Respiration predicts second-to-second variations of LFP sigma power

In the example of **Fig. 6A**, some bouts of pauses after inhalation (which corresponded to packet ends) coincided with sigma-band power variations, but were too subtle to be characterized as proper ‘troughs’. This suggests that changes of respiration modes are associated to variations of sigma-band power that are finer than the more drastic changes detectable as troughs. If so, we reasoned that we should be able to build an accurate predictor of sigma-band power based of respiratory features only. To test if this is true, we adapted the machine learning algorithm for state prediction (**Fig. 2**) by simply changing its output layer for z-score prediction and trained it for sigma-power regression during NREM (**Fig. 7A**) (**Methods**). To test the generalization of the respiration-LFP relationship across animals we again left one animal out of the training data set and used it for validation. In two example sessions we confirmed that the measured and predicted sigma powers fluctuated in the same direction in most instances, while the predictions of a control algorithm trained with shuffled data (**Methods**) remained flat (**Fig. 7B**). This translated into strong positive Pearson’s correlation between predicted and actual instantaneous sigma-power (0.58 ± 0.14, mean ± SD, N = 38 sessions), but not for the control algorithm (0.06 ± 0.13, N = 38 sessions) (Wilcoxon test, prediction *vs* shuffle: *P* = 7.74e-8, N = 38) (**Fig. 7C**). Moreover, the predicted sigma power was decreased at most sigma troughs in the examples (**Fig. 7B**). This was confirmed at the population level when plotting the measured and predicted sigma power around those events for all sessions: predicted sigma power was lowest at the time of the sigma trough (-0.91 ± 0.30 for the CNN predictions, mean ± SD, -0.11 ± 0.17 for the control; Wilcoxon signed-rank test *vs* shuffle: *P* = 1.35e-7, N = 38) (**Fig. 7D**). Finally, if most dips in predicted sigma power aligned with the end points of pause-after inhalation-based packets and/or sigma troughs in exemplar sessions, we also saw instances of co-varying predicted/actual sigma power outside of those (**Fig. 7B**, arrows), which showed that LFP and respiration can show subtle changes which are coordinated by general, cross-animal mechanisms.

**Figure 7.**
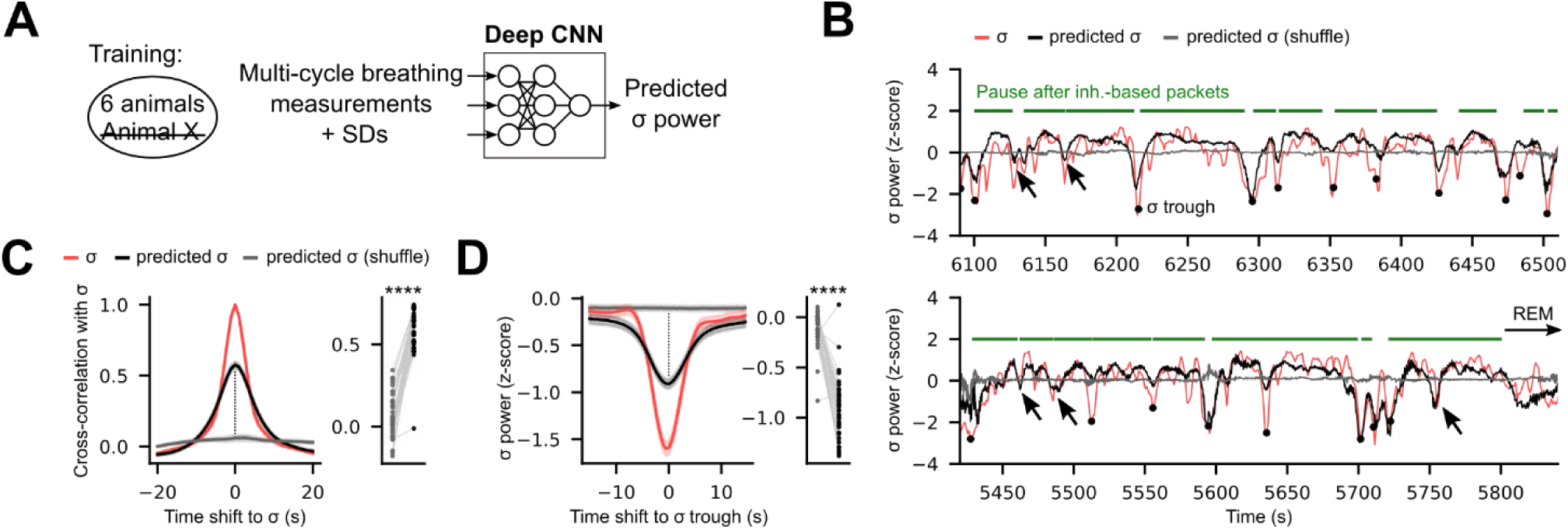
Respiration predicts sigma-band power fluctuations in NREM sleep. **A.** Machine-learning algorithm (convolutive neuronal network, CNN) for the regression of sigma-band power based on the 10 respiratory features (and their local SDs) (**Methods**). For validation, one animal is sequentially left-out the training data set. **B.** Example predictions of sigma-band power with the CNN algorithm for a left-out animal. Training with randomly shifted cycles (shuffle) as control (**Methods**). **C.** Cross-correlation between the measured sigma power and itself (auto-correlation), the predicted sigma power from respiration measurement, and the control predictions. Right-panel: correlation at shift 0 s with sigma power for the paired predictions of the CNN algorithm and the control algorithm across sessions. For the CNN prediction: 0.58 ± 0.14, shuffle: 0.06 ± 0.13 (mean ± SD). Wilcoxon test across sessions, prediction *vs* shuffle: *P* = 7.74e-8, N = 38 sessions. Z-score normalization was computed over NREM cycles. **D.** Sigma-band power around sigma troughs for the actual measurements, the respiration-based prediction and their control. Lines and shaded areas: mean ± SEM across 38 sessions. Lines and shaded areas: mean ± SEM across 34 sessions. Right-panel: sigma power at shift 0 s from the trough for the paired predictions from the CNN algorithm and the control algorithm. For the CNN prediction: -0.91 ± 0.30, shuffle: -0.11 ± 0.17 (mean ± SD), N = 38 sessions. Wilcoxon test across sessions, prediction *vs* shuffle: *P* 1.35e-7, N = 38 sessions.

## Discussion

In this study, we monitored hippocampal activity, head-acceleration and nasal pressure in freely moving mice equipped with portable pressure sensors. The precision of the pressure signal obtained revealed that mouse respiratory cycles not only comprise inhalation and exhalation, but also epochs of variable duration characterized by little or no airflow. The occurrence of pauses was specific for each brain state: in Wake, they mostly occur after inhalation, while they follow exhalation in REM sleep and were found in after both inhalation and exhalation during NREM. Pauses after inhalation varied independently from other features of respiratory cycles across states highlighting their unique nature. When predicting brain state based on cycle features with a GLM, we found that absence of pauses after inhalation contribute the most to the prediction precision for REM. In line with this observation, we found that transitions between vigilance states were accompanied - and sometimes preceded - by changes in respiratory pause duration, pointing towards a shared mechanism. We also demonstrated that respiratory pauses highlight the micro-architecture of NREM sleep by showing their coincidence with micro-arousals and even subtler troughs in sigma oscillation power. Based on the tight link between sigma power and pauses, we introduced new “respiration-based packets” delineated by bouts of pauses that capture NREM sigma-enriched time-windows. Overall, this work shows that pauses in nasal airflow are key components of the respiration cycles that contribute to the specific physiology of individual vigilance states and the finer architecture of sleep.

### Monitoring respiration in naturalistic conditions

Because of the central role of respiration for sensation and cognition (3, 5, 7, 8, 15, 43, 44), several methods have been developed over the years to monitor respiration in parallel with brain activity in animal models (42, 45–50). In head-fixed rodents pressure sensors connected to nasal cannula have allowed studying respiration-driven neuronal activity with high precision (33–36). Yet, measuring nasal pressure has remained challenging in freely moving mice, especially when it was combined with neuronal recordings (but see refs (7, 32, 51)). In these conditions, a commonly used method consists of measuring variations of nasal temperature using thermocouples or thermistors (47, 52–56). Alternatively, respiration can be monitored *via* whole-body plethysmography (57–59). Yet, these methods are not ideal when studying animals in freely moving conditions: thermocouples provide attenuated signals during sleep (in our conditions, the signal non-exploitable in this state, **Supplementary Fig. 1B**) and plethysmography restricts rodents to small hermetic enclosures and are thus incompatible with most naturalistic behaviors. Most importantly, both of these approaches do not allow a precise detection of fine respiratory features such as pauses that can only be revealed with accuracy through the use of pressure sensors. Respiration monitoring through pressure sensors has indeed proved useful when it comes to classifying wake behaviors (32). As respiration is a strong modulator of brain activity (2, 3, 44, 60) and has been proposed to act as a widespread synchronizer of neuronal networks (8, 15, 60), the methodology and analysis pipeline proposed here based on portable pressure sensors therefore represents a valuable technical advance for re-evaluating those results with improved respiration monitoring and in light of respiratory pauses.

### Scoring vigilance states with nasal pressure signals

Because the assessment of vigilance states is central to basic and clinical studies, a large numbers of automatic scoring methods have been developed, both for human and rodent models (reviewed in (38)). Most approaches rely on a combination of neuronal and movement signals (e.g. coming from electroencelography – EEG - electrodes and electromyograms - EMG) but one study proposed to use only local field potential signals from the hippocampus and the olfactory bulb (13). Given the strong link between respiration and neuronal activity in the olfactory bulb (4, 61–64) and the hippocampus (8, 55, 65, 66), the success of the latter approach is linked to the known changes in respiratory patterns across brain states (6–8, 14, 30). Another study showed that qualitative visual scoring of vigilance states based on whole-body plethysmography was in good agreement with manual scoring based on EEG and EMG data of individual mice (67). Yet, whether respiration changes are consistent across animals and whether these could be used, without additional information (such as EMG), to predict vigilance states was still unclear. Here we present a machine-learning-based method that provides automatic and accurate prediction of vigilance states based solely on nasal pressure signals (**Fig. 2**). Our results parallel those from a recent computer vision method for vigilance scoring which relied partly on visual features in videos which could reflect breathing (68). Video-based assessment of respiration was however limited to the rate and our precise characterization of the components of respiratory waveforms and their local variability, including pauses, can explain the superior performance of our approach for REM detection (**Fig. 2**). Moreover, we show that unlike amplitude features, the temporal properties of the respiratory components are comparable across animals (**Fig. 3**). This consistency likely allows our algorithm to provide an accurate prediction of vigilance states in new animals that were not part of the training set. The pre-trained network, thanks to its accuracy and generalization capability, could be applied to other datasets and become a valuable asset for state scoring based exclusively on respiratory pressure signals, without requirement for electrophysiological recordings.

### Respiratory features and neuromodulation throughout sleep stages

Our results provide strong evidence that the makeup of respiratory features significantly differ between NREM and REM stages of sleep (**Fig. 1**-**3**), with REM being strikingly devoid of pauses after inhalation. Moreover, local SDs for inhalation features were higher in REM as compared to NREM (**Supplementary Fig. 4B, dynamics at transitions in Supplementary Fig. 9**) consistent with previous studies showing erratic breathing in REM (9, 28). Local stability (as assessed by SDs) however showed only weak correlation with the amplitudes of the respiration parameters (**Supplementary Fig. 5C**) and may therefore reflect a parallel set of mechanisms linking respiration to brain states.

Looking a state transitions, we found that pauses after inhalation progressively shortened ∼100 cycles before animals entered REM sleep (**Supplementary Fig. 8I**). Previous studies linked these state transitions to variations in neuromodulator levels (69). For instance, acetylcholine levels in hippocampus are higher during REM as compared to NREM (22, 70–72) and during micro-arousals as compared to their neighboring NREM packets (22). Because we found opposite effects regarding inhalation pauses between REM and micro-arousals (a decrease in REM and an increase in micro-arousals), pauses after inhalation are unlikely to be functionally linked to the cholinergic system during sleep. Alternatively, norepinephrine (NE) released by the locus coeruleus (LC) has been shown to peak at micro-arousal occurrence during NREM, while its levels progressively decreases within NREM packets and when transitioning to REM sleep where they remain low (21, 73). NE dynamics therefore seem to mimic that of pauses after inhalation in NREM and REM sleep. Like NE levels (21), the durations of these pauses decreases within NREM packets and as sleep deepens, reaching their lowest values in the period that immediately precedes REM (**Fig. 5F**), also referred to as intermediate sleep (20, 21, 74, 75). We therefore hypothesize that the noradrenergic system is linked to the structure and physiology of respiratory pauses during sleep.

### Mechanisms of respiratory pauses

A wealth of studies addressing the neural basis of breathing has established a solid theoretical framework pointing at brainstem subnetworks as critical for setting the respiratory rhythm (1, 76–79). In particular, the pre-Bötzinger complex (pre-BötC) is considered crucial for driving inhalation whilst the retrotrapezoid nucleus/parafacial respiratory group is known to orchestrate active exhalation. Recent evidence showed that immediately after inhalation - an epoch termed “post-inspiratory phase of breathing” – a dedicated subnetwork of excitatory neurons with autonomous rhythmic properties termed post-inspiratory complex (PiCo) is recruited (80). Based on this discovery, the “triple oscillator” hypothesis has been proposed where the three phases of breathing are governed by distinct brainstem subnetworks (78, 79). In this framework, our report of respiratory pauses occurring after inhalation seem consistent with the concept of the post-inspiratory phase, and it is plausible that PiCo neurons are involved in driving pauses. Yet, the fact that pauses show variable durations partially challenges the view that respiration results from a pure oscillatory phenomenon. Based on our results, we propose instead that mice can block outgoing exhalation airflow after inhaling, and that this blockade dominates during wakefulness in freely-moving conditions. The neural mechanisms at play during these pauses have to be further studied in light of sensitive pressure sensor recordings. Yet, as explained above, the progressive elimination of pauses after inhalation through NREM sleep (**Fig. 5E, F**), until complete elimination in REM, already prompts us to propose that the noradrenergic system, but not cholinergic one, participates in the control of pauses after inhalation. In support of this hypothesis, LC is involved in the control of breathing (81, 82) and sends information to the medullary respiratory network (83, 84). In an alternative model to consider, fluctuations in NE concentrations would result from changes in respiratory behavior since LC receives inputs from the medullary respiratory network (85) and shows respiration-modulated activity (86). Future work is required to unveil the modalities of the interactions between neuromodulatory systems and respiration centers at the transitions between sleep sub-stages.

As for pauses after exhalation, they rarely occurred in awake, freely moving animals, but were prominent in NREM and dominated REM sleep. In fact, their dynamic generally corresponded to a mirrored image of the dynamic adopted by pauses after inhalation during sleep stages. Because REM sleep is characterized by muscular atonia we suspect that the inflections detected in the pressure signal (that defines pause after exhalation onset) is related to the physical properties of the lungs that passively deflate following inhalation. Interestingly, central sleep apneas observed in humans (87) and mice (88) occur at very low rate (∼22 apneas/hr in mice with the most permissive detection criteria (89)) as compared to the pauses we study here (present in nearly all breathing cycles). Yet, apnea events have been shown to occur at the end of exhalation, similar to the pauses we identified during sleep. It raises the possibility that central sleep apneas, which affect sleep quality and are linked with cardiovascular dysfunctions (90–92), share similar mechanisms with the physiological respiratory pauses highlighted in our study. Clarifying the neuronal substrates involved in sleep and respiration regulation could therefore improve our understanding of the mechanisms underlying detrimental breathing disorders in sleep.

### Respiration reveals the micro-architecture of NREM sleep

The micro-architecture of sleep and NREM packets are currently raising attention as they orchestrate homeostasis (20) and memory (21) processes in sleep. Specifically, spindle-rich epochs are nested at the center of those packets indicating that they could determine specific time-windows for system-level memory consolidation (19, 26, 27).

Despite their importance, the definition of NREM packets differs across studies. Most of them use movement-related signals and drops in LFP delta power to detect micro-arousals and therefore packet boundaries (20, 22). Our study indicates that respiration - and in particular respiration pauses – abruptly and consistently changes around micro-arousals associated with movement but also during NREM periods when no movements are detected. This echoes a recent study which suggested that fluctuations in NE levels can reveal micro-arousals in absence of detectable movement changes (21). We show that these transient respiratory changes tightly track the intermittent drops in LFP delta and sigma power suggesting an increased sensitivity to micro-arousals. In line with previous descriptions of infra-slow oscillations (23), our data show that the packets outlined by respiration changes are more homogeneous in durations and in LFP power than packets defined by movements. Based on these observations, we think that “respiration packets” constitute an ideal time unit when it comes to establishing NREM micro-architecture. Finally, while recent work highlighted the link between the late phase of breathing cycles and spindle occurrence in the human brain (93), our work focuses on longer timescales and shows that not all cycles are equivalent across NREM sleep (notably with regards to pauses): the presence of late pauses following exhalation may set the stage for spindles and memory processes at the end of NREM sleep packets.

## Methods

### Animals

All experimental procedures were performed in accordance with standard ethical guidelines (European Communities Directive 86/60-EEC) and approved by the local committee on animal health and care of Bordeaux and the French ministry of agriculture and forestry (authorization numbers 18625, 19746, 23974 / facility agreements A33063940 and A33 063 943). All mice were maintained in pathogen free facilities in a diurnal 12h light/dark cycle with food *ad libitum*. A total of N = 21 mice were used in this study: 8 animals (N males = 7, N females = 1) were implanted with nasal cannulas and hippocampal silicone probes, 12 male mice were used for odor detection task for behavioral validation (N = 7 sham, N = 5 Cannula), 1 male mouse was implanted with intranasal cannula and thermocouple for methodological validation. All mice were C57bl6/J except for one OXT-IRES-cre female mice.

### Pressure sensor

Pressure sensors (PS) were purchased from Honeywell (part #: SSC S RN N 004ND AA5). Our protocol is based on a previous approach (33) that we adapted to recordings in freely moving mice in two ways. First we added a 6 pin male connector below the PS to fit a 6 pin female connector cemented on the head of the animal. This ensured a stable fixation of the PS during recordings in freely moving mice. Second, we soldered the input and output electric connections of the PS to thin Litz wires which were themselves held by a pulley system ensuring very low weight on the head of the animal. A piece of polyethylene tubing (801000, A-M Sys-tems, ID 0.015in, OD 0.043 in) of adjustable length allowed to connect the PS port to the nasal cannula. During inhalations, the inward flow of air into the nose causes a decrease in the measured pressure. During exhalations, the outward flow of air from the nose results in an increase in the measured pressure. Besides these inhalation- or exhalations-induced deflections, flat pressure signal corresponds to atmospheric pressure and respiratory pauses.

### Nasal cannula

Nasal cannulas were home made from 23G hypodermic stainless steel tube (A-M Systems). Briefly, the tube was cut to an 8mm length and the tip was beveled with a 45 degrees’ angle. A mark at 1.5 mm from the beveled tip symbolizes the limit for insertion during surgery. To ensure that cannulas do not get blocked by dust or litter between recording sessions, they are capped with dummy plugs when animals are not being recorded. Dummies consist in 11mm long 27Gx1/2’’ industrial dispensing tip (CML supply). Only 2mm of plastic is kept from the top of the dispensing tip to allow for an easy removal by the experimenter. On this dummy a 3mm piece of 23G stainless steel tube and an 8mm of polyethylene tubing (801000, A-M Systems, ID 0.015in, OD 0.043 in) are glued to the base. The 3mm tube acts as a stopper to ensure that the dummy does not exceed the length of the cannula, and the polyethylene tubing holds the dummy onto the cannula.

### Thermocouples

Thermocouple-based measurements of respiratory behaviors rely on the fact that the body temperature of mice (38℃) is warmer than external temperature in our recording conditions (20-24℃). During inhalations, the inward flow of external air into the nose causes a decrease in the measured temperature. During exhalations, the outward flow of air from the nose results in an increase in the measured temperature. Here we used K-type thermocouples (Omega) inserted in the nasal cavity of mice and cemented in place on the skull (see below).

### Surgery

Surgeries for local cannula implantation were performed as previously (33, 34). For local cannula implantation, mice were anesthetized using isoflurane (induction 3 minutes at 4%, then at 1.5% during the surgery). They received an intraperitoneal injection of an analgesic (Metacam, 5mg/kg) and hair above the skull was removed using a hair removing cream (Veet). Mice were then placed on a stereotaxic frame (Kopf) where vitals (body temperature, heart rate, blood oxygenation level) were tracked using PhysioSuite (Kent Scientific). The eyes were protected against dryness with Vaseline. A midline incision was performed above the skull and the nasal bone following a local subcutaneous injection of 0.1ml of lurocaine (5mg/kg). The skull was cleaned of any conjunctive tissue using a micro-curette (Fine Scientific Tools). A dental drill (small size) was used to perform a craniotomy in the nasal bone. A single and fast movement of the drill ensures effective opening of the nasal epithelium membrane located right below the nasal bone. During this delicate step, it is important not to touch the turbinates located below the hole with the drill tip to prevent any subsequent clogging of the cannula. Coordinates for optimal respiratory recordings (AP = 3.5-4, ML = 0.5, DV = 1.5mm) were defined from the junction between the midline and the fronto-nasal sutures (but these can potentially vary depending on the mouse strain and age). During insertion, the beveled tip of the cannula was oriented towards the midline to increase successful signal collection. Cannula was implanted with its dummy to avoid any blood or tissue to enter the cannula during insertion. Nasal cannula is then fixed to the bone via Superbond (C&B), and a 6 pin connector is cemented on the back of the skull to allow attachment of the pressure sensor in subsequent recordings (see <u>Pressure sensor</u> section). Finally, a handle was cemented on the skull to facilitate the immobilization of the mouse during future pressure sensor attachment procedures and the skin is closed with Vetbond (3M). Sham animals used in the odor detection task (**Supplementary Fig. 1**) underwent the same surgical procedure except that no craniotomy was performed and the cannula was cemented at the surface of the skull, above the nasal bone.

For 8 mice, high density silicon probes (NeuroNexus) were implanted in the CA1 region of the hippocampus (AP = -1.8, ML = 1.4, DV = -1.2). The local field potentials recorded in this region combined with the accelerometer signal obtained from the Intan headstages (RHD 32ch, Intan Technologies), allowed brain-state annotations.

For the comparison of the signals obtained from PS and thermocouple (K-type TC, Omega), two symmetrical craniotomies were performed above the nasal bone at the same AP coordinates as for PS implants. PS was implanted as described above. For TC, removal of the nasal bone was performed by progressively thinning the skull with gentle drilling until the highly irrigated nasal epithelium membrane was revealed. The membrane was pierced with a fine cotton tip, creating the hole required to insert the TC 1.5mm deep into the cavity. The craniotomy was then protected with Kwik-Sil (WPI), and the implant was stabilized with Superbond (C&B). For brain state detection, electroencephalogram (EEG) signals were collected using a miniature screw implanted in the cortical bone and referenced to a ground screw above the cerebellum.

### Plethysmograph

Whole-body plethysmography (57–59) provides a non-invasive biomechanical measure of respiration. Here, a whole-body plethysmograph chamber (Emka Technologies, France) was used to record mouse respiratory activity in parallel with nasal pressure monitoring. Mouse breathing induces pressure changes in the plethysmograph chamber. These were captured by a differential pressure sensor that compares pressure in the animal chamber with the reference chamber. Constant airflow (2.2 L/min) was provided through the apparatus. The respiratory signal collected from the plethysmograph was interfaced to a computer equipped an Intan acquisition board (see Data acquisition section).

### Data acquisition

Pressure sensor, thermocouple and plethysmograph signals were acquired continuously at 20 kHz on an Intan RHD2000 interface board analog input channels (Intan Technologies). A voltage divider was placed between the pressure sensor and the acquisition board to insure that the voltage range was not exceeding 3.3V. Thermocouple signal was amplified before acquisition (amplification factor: 192). Electrophysiological signals were simultaneously acquired at 20 kHz after being amplified by 32 and 64-channel digital headstages (Intan Technologies).

### Brain state scoring

Since past studies have demonstrated that the presence of hippocampal theta oscillation in an animal with sleeping posture is sufficient to identify Rapid eye movement sleep (REM) sleep (94), epochs of Wake, REM and non-REM (NREM) have previously been defined using the ratio of the hippocampal LFP power in theta band (5–11 Hz) and delta band (1–4 Hz) combined with motion signal (39, 95). Similarly in our study, Wake, REM and NREM episodes were annotated offline using the ratio of the hippocampal LFP power in theta band (5–11 Hz) and delta band (1–4 Hz), the accelerometer signal as well as visual inspection of raw traces and whitened power spectra (using a low-order autoregressive model), but with the help of a Matlab GUI (see *TheStateEditor* from the BuzsákiLab repository, https://github.com/buzsakilab/buzcode/blob/dev/GUITools/TheStateEditor/TheStateEditor.m) (40). Several previous studies have demonstrated that the presence of hippocampal theta oscillation in an animal with sleeping posture is sufficient to identify REM sleep (94).

To label breathing cycles used to train the artificial network, we used an automatic state scoring approach described previously that relies on a combination of CA1 LFP spectral features and motion signal (41) (StateScoreMaster from the Buzsáki Lab (https://github.com/buzsakilab/buzcode/blob/dev/detectors/detectStates/SleepScoreMaster/SleepScoreMaster.m), and adapted it to account for the heterogeneity in mouse behavior across sessions (e.g. variability in sleep duration between sessions/mice) following these steps:

1. In each individual session we obtained time-series data (time bin = 1 sec) of:

a. EMG as the correlation between electrophysiological signals recorded from channels at the ends of each shank filtered in the 300-600Hz band (see Schomburg et al., 2014).
b. slow-wave ratio as the slope of the CA1 LFP power spectrum (4-90 Hz) (see Gao, et al., 2017, Watson, Dingm Buzsaki 2018) ;
c. theta ratio as the peak of the slope CA1 LFP power spectrum in the theta band (5-10 Hz) (see Gao, et al., 2017, Watson, Dingm Buzsaki 2018);
2. The EMG, slow-wave and theta ratio obtained in each session were normalized (0–1), and then pooled across all sessions revealing overall bimodal distributions.
3. Population threshold indicating boundaries between low to high EMG, slow-wave and theta-ratio were determined as the values showing peaks of the inverted distributions: EMG threshold = 0.38, slow-wave ratio threshold = 0.46; theta ratio threshold = 0.46.
4. In each session, state intervals were then computed using population thresholds: wake (high theta-ratio, high EMG), NREM state (high slow-wave ratio, low EMG) and REM state (high theta-ratio, low EMG).
5. For the automated sleep scoring two extra conditions were applied: a) intervals assigned as REM following wake intervals (which is known to be an impossible condition) were corrected as NREM; b) NREM intervals with duration <30 seconds surrounded by two wake intervals were corrected as wake.

Visual examination of the results obtained from automated sleep scoring compared to that from manual annotation were largely consistent across sessions, justifying their use for ground truth values for the brain-state prediction algorithm.

### Pressure sensor data analysis

All analyses were performed using MATLAB (The MathWorks) built-in functions, the FMAToolbox (http://fmatoolbox.sourceforge.net/), Buzsaki lab toolbox – *buzcode* (https://github.com/buzsakilab/buzcode), the freely available MATLAB toolbox from the Zelano lab *BreathMetrics* (https://github.com/zelanolab/breathmetrics) and custom-written scripts.

Respiration was analyzed *post-hoc* using a modified version of BreathMetrics, a toolbox designed to automatically describe respiratory features from pressure sensor signals acquired from human or rodent subjects (31). We called this modified version “mouse-BreathMetrics” to acknowledge the contribution of the original script and highlight the fact that our script was only tested on mouse data. Here, we used *BreathMetrics* with a redundant re-iterative approach aimed at maximizing the number of detected respiratory cycles. This was done in order to both account for signal instability of our recordings (baseline drift, electrical noise, transient loss of signal) and to prevent potential artefacts due to radical changes in the respiration pattern across behavioral states. The steps followed to analyze respiratory signal of each session are summarized here and can be visualized in **Supplementary Fig. 2 and Supplementary Movies 1-3**.

Firstly, the respiratory signal was examined by the experimenter to ensure that only intervals with valid data, “valid sniff-blocks”, were taken into account (ex 0-20s, 40-80s, 95-160 etc in **Supplementary Fig. 2 Ai**). This step was only necessary for sessions with transient nasal cannula clogging that results in pressure signal loss.

Secondly, valid sniff-blocks were intercepted with brain states (see **Supplementary Fig. 2 Aii Wake**: 0-60 seconds, 200-240 seconds, 280-300 seconds; NREM: 60-140 seconds, 160-200 seconds, 140-260 seconds, 240-255 seconds, 260-280 seconds; REM: 140-160 seconds, 255-260 seconds) previously detected using hippocampal LFP spectral features (theta/delta ratio) and accelerometer (movement) as previously described (see **Brain state scoring** section of **Methods**)(40). The resulting “sniff-chunks” contained homogeneous signal belonging to only one behavioral brain state (see **Supplementary Fig. 2 Aiii**; Wake chunk: 0-20 seconds, 40-60 seconds etc; NREM chunk: 60-80 seconds, 95-140 seconds; REM chunk: 140-160 seconds, 255-260 seconds).

Thirdly, each sniff-chunk was individually analyzed using *BreathMetrics (***Supplementary Fig. 2 Bi**), so that a preliminary characterization of the respiratory cycles could be obtained (**Supplementary Fig. 2 Bii**). However, we noticed that errors in the detection of the onset/offset of the respiratory components were frequently encountered mostly due to the inaccurate pause detection. More rarely, respiratory cycles were found to be “skipped” due to short epochs of baseline-drift in the raw signal, heavily altering the quality of the results during those short windows. To account for this, we used BreathMetrics with a redundant re-iterative approach to maximize the number of respiratory cycles in each sniff-chunk. We further split each sniff-chunk in “sniff-lets”, short time intervals containing the signal of 10+1 pre-detected respiration cycles (see **Supplementary Fig. 2 Ci**). Theoretically each sniff-let contained 11 cycles but this was ensured by iteratively altering its signal with a moving-average window correction of increasing size (20 iterations, window size 1-3 seconds). In each iteration, the altered signal was run with *findExtrema* from *BreathMetrics* toolbox yielding each time the number of detected respiratory cycles. At the end of 20 iterations, the altered signal with highest number of sniff cycles was chosen for the next steps – otherwise the raw signal was maintained. Once ensured each sniff-let contained the highest number of cycles, the signal was re-examined using the following functions from *BreathMetrics* toolbox:

Breathmetrics
correctRespirationToBaseline
findExtrema
findOnsetsAndPauses
findInhaleAndExhaleOffsets
findBreathAndPauseDurations
findInhaleAndExhaleVolumes

In order to obtain a finer detection of respiratory components (in particular pauses during wake state), we took into account minima and maxima of the sniff-let raw signal first derivative reflecting troughs and peaks in the rate of instantaneous pressure (see **Supplementary Fig. 2 Cii**).

In each sniff-let, inhalation and exhalation were re-calculated after pause detection which were assigned if the following criteria were simultaneously met (see **Supplementary Fig. 2 Ciii**):

a. pressure signal in the 5-95^th^ percentile range during previously assigned pauses epochs of the entire sniff-chunk;
b. first derivative in the mean + standard deviation of the first derivative signal during previously assigned pauses epochs of the entire sniff-chunk;
c. outside of the window between ascending peak and descending through of the first derivative of the sniff-let (putative inhalation);
d. outside of the window between descending through and ascending peak of the first derivative of the sniff-let.

### Single-cycle data normalization and analysis

In **Fig. 2-4**, we analyzed single-respiratory-cycle properties of recordings from 7 animals (males and females): 1) id 3C028 (3 sessions), 2) 3C030 (14 sessions), 3) 3C060

(7 sessions), 4) 3C209 (2 sessions), 5) 7C012 (6 sessions), 6) 7C014 (5 sessions), 7) 7C026 (5 sessions). Brain states were annotated using the method described above with CA1 LFP signals that were simultaneously recorded and accelerometer data.

For the machine learning-based approach and state transition analysis (**Fig. 2-4**), we ran BreathMetrics on each valid sniff-block, without breaking out the signal into state-specific chunks, hence making cycle detection and feature extraction state-blind. The same analysis pipeline as the one described above was conducted. Respiratory features extracted by *BreathMetrics* were: peak amplitudes, volumes, peak times and durations for both inhalation and exhalation phases, as well as durations of pauses after inhalation and exhalation. To allow for principal component analysis (PCA) and state prediction each feature values were first log-normalized to make their distribution Gaussian-like (x’=*log (x + 10e-4)*). Then, each log-feature was centered with respect to the median of each animal mean value (m) (*n* = 7) (computed over all its cycles) and normalized by the median of each animal standard deviation value (s). Normalization parameters (**Supplementary Table 1**) were fixed *a priori,* and were used unchanged for all sessions from all animals. Doing so, the standard deviation of each feature was approximately (but not strictly) 1 for each session and each mean was approximately 0.

For automated brain state prediction with a deep CNN (**Fig. 2D-I**) log-features were augmented by their temporal standard deviation (SD) over a local time-window of size 10 cycles. Those SDs were themselves centered and unit-variance normalized (**Supplementary Table 1**). In this case we used automated state annotation as provided by the method described above so as to avoid expert-to-expert differences and inconsistencies which cannot be learnt.

Principal component analysis of single cycle variations (**Fig. 3** and **Supplementary Fig. 6**) was performed by singular value decomposition of the covariance matrix of the normalized features. The latter matrix was obtained by equally sampling cycles in sessions from all animals (2000 cycles for each animal-brain state pair). Entropy values in **Fig. 3D** were computed by building a histogram of PC coefficient values with 200 equally spaced bins and then computing for each bin the empirical probability p_j_ = *Pr(animal id|coefficient value in bin #j)* based on the proportion of animals associated to the coefficient in the bin #j. The entropy of this bin was computed as -*p_j_log(*p_j_) and the total entropy was the weighted sum of the entropy values of all bins, with the proportion of coefficients falling in a bin as its weight. The same procedure was applied for entropy values in **Fig. 3F**, but the empirical probability p_j_ = *Pr(brain state|coefficient value in bin #j)* was computed and used instead.

### Automated brain state prediction from respiratory features

Automated brain sate prediction based on a generalized linear (GLM) model was achieved by using the 10 normalized features from single respiratory cycles as inputs. Linear outputs of the 3-states model were compared to automated state predictions based on LFP signals and GLM parameters were adjusted to minimize their cross-entropy. For GLM training, 1e4 respiratory cycles were randomly drawn for each animal, and 80% of those were used for parameter adjustment, while the remaining 20% were used for validation and performance computation as shown in **Fig. 2B-C**. All fitting procedures were repeated 10 times to characterize performance variability. For the *+Local* SDs condition, input parameters were augmented by their local standard deviations, as described above. The weights of the Wake/NREM/REM classes in the objective function were set to the inverse of their occurrence rate to equalize their importance (balanced condition used in **Fig. 2B**). For computing receiving operating characteristics (**Fig. 2C**), the weight of the REM state was changed stepwise to yield varying recall *vs* false positive rates results. The GLMs were implemented in Python and fitted to the data by using the sklearn module.

The design of our artificial network specialized for brain state prediction is shown in **Supplementary Fig. 6**. It takes 201 breathing cycles described by the 20 normalized features (including local SDs) as inputs and the output layer combines both local (first convolutive layers) and contextual information (last convolutive layers) to compute the probabilities of each three states.

As a final step, state probabilities are smoothed using a temporal Gaussian filter (with σ = 20 cycles) and the predicted state is taken as the one with maximal probability. The confidence in the prediction is computed as this predicted state probability so that confidence is high when the differences between state probabilities are large.

The algorithm was implemented using a Jupyter notebook with Python and is made freely available. Numpy and Scipy libraries are mainly used for mathematical computation. The proposed artificial neuronal network structures are implemented thanks to the TensorFlow (2.0.0) dedicated library with GPU support. 2733 free parameters need to be adjusted for best prediction performance.

Network parameter optimization was carried out using sessions from 6 animals, leaving one out for assessing data overfitting and for validating the ability of the network to infer generalized properties of breathing features and brain states to other animals. Training was repeated 6 times, cycling through different animals for validation. Parameter optimization was performed by minimizing the categorical cross-entropy between the discrete annotations of states (ground-truth) and the continuous prediction of their probabilities by the network. To cope for the imbalance between the frequency of each state (REM being the rarest), different weights were used for the prediction of sates: 0.1 for wake cycles, 0.25 for NREM, and 0.5 for REM. A gradient-descent-type algorithm (Nadam) was used with learning rate 10e-5. We re-sampled 25 times (outer epochs) 2000 random cycles from each animal (all sessions) to limit memory usage. 3 steps of gradient-descent were performed with those 2000×6 = 12000 cycles (inner iterations). For training only, ‘dropout’ (with a rate of 10%) layers were inserted between the layers to regularize the optimization problem and thus limit data overfitting. 10 independent networks were trained for results averaging in **Fig. 2F-I**.

### Analysis of breathing dynamics at state transition

For each annotated state transition, we computed PC1 and 3 coefficients for the 300 respiratory cycles right before, and the 300 coefficients right after, the transition time point. Individual ± 300 cycles series were clipped so that extra state transitions could not occur around the transition of interest. All transitions from one type were time-aligned and averaged to obtain mean curves in **Fig 4. G, H**. We computed the amplitude change as the difference between averaged coefficient values post and prior to state transition over those time-windows. The statistical significance of each change was computed using a KS test between values posterior and prior to state transition. When the change was significant (*P <* 0.05) we fitted a shifted and scaled logistic function to coefficients: *PC value(t) = amplitude/(1 + exp(-speed x (t – t_0_)) + baseline* if the mean trace (**Fig 4. G, H**) was increasing, and *PC value(t) = amplitude x (1 - 1/(1 + exp(-speed x (t – t_0_))) + baseline* if it was decreasing. *t_0_* = 0 would correspond to a sigmoid function time-centered to the state transition. We used constrained optimization (MATLAB’s *fmincon* function) for parameter fitting and we kept only values for fitting procedures that succeeded in **Fig. E, F**. In **Fig. 4D**, amplitude values resulted either from the fitting procedure when available, or from the simple pre-post difference (as described above) otherwise. d_onset_ (time to 10% amplitude change) and d_offset_ (time to 90% amplitude change) were computed as: *d_onset_ = -log(1/0.1 - 1)/amplitude + t_0_* and *d_offset_ = -log(1/0.9 - 1)/amplitude + t_0_*.

### Microarousals detection and associated NREM packets

To detect micro-arousal events occurring during NREM intervals of sleep within each individual session, we adapted methods from previous study (42) as follows:

a. all three accelerometer signals (sampling rate = 20KHz) were mean-subtracted and then summed into a single one;
b. local standard deviation of the acc. signal was obtained by first computing the local variance (5 second window) of the summed acceleration signal and then square rooting it;
c. motion threshold for putative microarousal events was calculated as: threshold = median(x) + k * 1.4826 * MAD(x)

where: x = local standard deviation of acc. signal during episodes of NREM sleep (first/last 30 seconds excluded), k = 3, MAD = median absolute deviation computed as median(abs(x-median(x))).

Micro-arousals were detected as those intervals occurring during NREM sleep episodes exceeding the motion threshold, lasting at least 300 ms and whose peaks were less than 5 seconds spaced (otherwise they were merged as a single event).

Finally, each NREM interval was fragmented into chunks surrounded by micro-arousals events. Only those chunks with NREM intervals before and after the micro-arousals were considered as valid NREM packets, whose duration was then normalized between 0 (start) and 1 (end) as in (22).

### Respiratory features during NREM packets

To address modulation of respiration during NREM packets, all the respiratory features of distinct respiratory cycles were first converted into time-series data by averaging the features of cycles falling within each time bin (size = 200 ms) during the entire session.

### LFP spectral analysis during NREM packets

To address hippocampal LFP changes during NREM packets, the hippocampal channel showing largest ripple power (4**th** order Butterworth filter= 140-180 Hz) during NREM sleep in each session was examined. Raw LFP sampled as 20KHz was low-pass filtered (450 Hz cut-of band) and then downsampled at 1250 Hz to compute Chronux multi-taper spectrogram (window = 5 seconds, step = 1 seconds, 0.5-25 Hz frequency range). Instantaneous delta/sigma power during NREM was calculated as the peak power in the corresponding frequency bands (delta = 0.5-4 Hz, sigma = 10-20 Hz) across time, smoothed with Gaussian kernel (sigma = 2 seconds) and z-scored after REM and wake interval exclusion. Delta/sigma troughs were identified as the local minima in the delta/sigma power with z-score <= 1 and minimum spacing of 5 seconds between consecutive events.

### Detection of respiration-based NREM packets

Detection of NREM packets based on pauses after inhalation was carried out as follows: 1) the duration of pauses after inhalation was averaged over 2 s windows and sampled every 0.5 s, 2) packets endpoints were delineated by bouts of long pauses (duration of pauses after inhalation > 50 ms for a duration ≥1.5 s), 3) only packets between those bouts that lasted at least 5 s were considered. This procedure resulted in NREM packets with homogeneously low pauses after inhalation (**Fig. 6E**).

### Automated prediction of the power of the LFP sigma-band from respiratory features

Prediction of instantaneous sigma power was based on a similar CNN as above, except for the output layer which consisted in a single unit with linear activation function to allow for z-score prediction. This resulted in network with 3178 trainable parameters. As an objective, we now minimized the mean squared error between the prediction and the z-scores of the sigma-band LFP power at NREM cycles only. We used 35 outer epochs for cycle resampling and kept the training rate at 1e-5. Again, we sequentially left one animal out for validation, resulting in 8 distinct CNNs (one for each validation animal), and all results were obtained by predicting sigma power with the network corresponding to the left-out animal. Unlike for state prediction (**Fig. 2**), since we focused on NREM solely, we used respiration features obtained from state-specific BreathMetrics which are most accurate.

### Explorative odor detection task

The explorative odor detection task conducted as previously (37). Briefly, the materials used consisted of a test cage similar to the home cage but with an odor holder that was made from a stainless-steel lid of an unused mouse water bottle. The bottle lid fit flush in a hole which prevented it from being displaced by the mice during the test. The tip of the lid’s nozzle extended 5cm and contained a 3mm hole from which odors could emanate. Cameras were installed above the test cage and on each side of the odor holder to allow easier visualization. The cage was placed on a table in a dedicated testing room (separate to the housing room).

The behavioral protocol consisted of 2 days of habituation to the test cage before the test day. Each mouse was placed in the test cage for 3 minutes each of 5 separate trials for the odor detection experiment with a 3-minutes inter-trial interval. Before each trial, 10 μL of mineral oil was placed on a 2cm strip of filter paper and placed in the odor holder. All mice were water deprived after the 2^nd^ day of habituation for 24 hours before the test to increase their motivation to explore.

Based on a serial dilution method with mineral oil, increasing concentrations of the odor (Isoamyl acetate [banana-like (Sigma-Aldrich)]) was tested as follow: mineral oil, 0.001%, 0.01%, 0.1% and 1%. 10μL of these solutions were placed on a strip of filter paper immediately before each trial and then introduced inside the nozzle of the odor holders. The odor holders were cleaned with 4% sodium bicarbonate and water between trials and experiments.

The time mice spent investigating the presented odor was counted manually using a customized program (BehavScor v3.0 beta): considered epochs corresponded to the time intervals when mice directed their nose <1cm from the tip of the holder. Mice exploring less than a total of 5 seconds within the 5 trials were excluded from the analysis and each animal was tested only once with a single odor. The odor threshold was defined as the odor concentration the most explored (relative to odorless mineral oil) among the 4 odor concentrations.

### Statistical Analyses

Statistical analyses were performed using scripts in MATLAB and Python (numpy and scipy libraries). All the results comparing respiratory features between vigilance states were performed after extracting the median values for each state in all sessions and compared using Kruskal Wallis test followed by *post-hoc* paired Wilcoxon signed-rank tests in the two-tailed configuration with Bonferroni corrections, Tukey’s honest significant difference procedure or Dunnett’s tests to account for multiple comparisons, as stated. Pooled distributions were obtained by randomly selecting 500 cycles for each state across sessions. In the 3 sessions where the number of REM cycles was <500, the number of selected cycles for Wake and NREM was matched to the number of cycles in REM. Overall we pooled N = 18107 cycles in each brain states. Additional nonparametric tests (Mann-Whitney U tests, Kolmogorov-Smirnov tests or Wilcoxon signed-rank tests in the two-tailed configuration) were conducted as stated in legends. Linear regressions were conducted using Pearson’s linear correlations and tested using a Student’s t distribution (MATLAB *corr* function). Significance was set with alpha = 0.05 and was represented on graphs as the following: * = *P <* 0.05, ** = *P <* 0.01, ***= *P <* 0.001, **** = *P <* 0.0001. As mentioned in the figure legends, individual data-points are plotted above bars indicating mean ± SEM. Box plots represent Q1 (25^prctile^), median and Q3 (75^prcile^) of the distribution, lower/upper whiskers show the adjacent values calculated with W = 1.5: lower value = Q1 - W*(Q3-Q1); upper value = Q3 + W*(Q3-Q1).

## Acknowledgments

We warmly thank Dmitry Rinberg and Christopher Wilson for sharing methodological details regarding nasal cannula implants. We thank the Zelano lab past and current members of the Buzsáki Lab, for making the Breathmetrics Toolbox and other scripts available. We thank Sophie Bagur and Karim Benchenane for their useful comments on the early version of the manuscript. We thank the Animal Facility of the Broca Institute (Pôle In Vivo) for mouse care and the Roux lab members for useful discussions. This work was supported by CNRS (L.R., F.G., E.H.), INSERM (N.C.), the Ministère de l’Enseignement Supérieur et de la Recherche (to C.M.), European Research Council (ERC-StG-851560 to L.R.), Bordeaux University (2017 IdEx Junior Chair ANR-10-IDEX-03-02 to L.R.), Fondation pour la Recherche Médicale and Fondation Schlumberger pour l’Education et la Recherche (FSER202112014572 to L.R.).

## Contributions

A. C., N.C., L.R. designed research; G.T., C.M, G.T., P.R., C.M. performed recordings in freely-moving mice; C.M., E.L. refined nasal pressure recordings; C.M. and D.J. performed recordings with plethysmograph; T.D. performed behavioral tasks; G.C. and N.C. perform the analyses; E.H., F.G., N.C. and L.R. supervised research; G.C., N.C. and L.R. wrote the paper and E.H. edited it. All authors approved the manuscript.

