## Supplemental Information for "Respiratory pauses highlight sleep architecture in mice"

### Supplementary tables

**Supplementary Table 1: Parameters for the normalization f(x) = ((x + 10e-4)-m)/s) of respiratory log-features extracted by BreathMetrics.**

| Log-feature | m | s |
|---|---|---|
| Inhalation amplitude | 0.37083446 | 0.42488484 |
| Exhalation amplitude | 0.32613887 | 0.63292769 |
| Inh peak time | -3.52245613 | 0.81363155 |
| Exh peak time | -3.71579229 | 0.62328763 |
| Inh volume | 3.92810079 | 0.40672072 |
| Exh volume | 4.0691885 | 0.47766716 |
| Inh duration | -2.90744862 | 0.61724445 |
| Exh duration | -2.72445354 | 0.59518787 |
| Pause after inh duration | -3.56154574 | 2.0797817 |
| Pause after exh duration | -8.1313956 | 2.15578837 |
| SD of Inh amplitude | 0.15100771 | 0.09135163 |
| SD of Exh amplitude | 0.22566782 | 0.12422676 |
| SD of Inh peak time | 0.25739959 | 0.17761934 |
| SD of Exh peak time | 0.33116313 | 0.2267266 |
| SD of Inh volume | 0.2200343 | 0.14223361 |
| SD of Exh volume | 0.30550223 | 0.18434577 |
| SD of Inh duration | 0.18682609 | 0.14933253 |
| SD of Exh duration | 0.23988165 | 0.15500514 |
| SD of Pause after inh duration | 0.97396974 | 0.69089047 |
| SD of Pause after exh duration | 1.17077256 | 1.09730831 |

**Supplementary Table 2: performance values for parameter exclusion conditions (mean±SD) (Fig. 2B)**

a) Recall (N = 10 for each condition)

| | **Wake** | **NREM** | **REM** |
|---|---|---|---|
| **All measurements** | 0.9163±0.0029 | 0.7605±0.006697 | 0.8427±0.0216 |
| **Shuffle** | 0.5174±0.0283 | 0.2469±0.0315 | 0.3803±0.0246 |
| **- Volumes & flows** | 0.9101±0.0040 | 0.6817±0.0076 | 0.8416±0.0166 |
| **- Inh & exh kinetics** | 0.8834±0.0046 | 0.7465±0.0071 | 0.8389±0.0211 |
| **- Pauses after inh** | 0.9023±0.0025 | 0.7358±0.0094 | 0.7323±0.0178 |
| **-Pauses after exh** | 0.9155±0.0032 | 0.7688±0.0099 | 0.8540±0.0179 |
| **+ Local SDs** | 0.9284±0.0023 | 0.8344±0.0107 | 0.8791±0.0148 |

b) Precision (N = 10 for each condition)

| | **Wake** | **NREM** | **REM** |
|---|---|---|---|
| **All measurements** | 0.9613±0.0016 | 0.8485±0.0044 | 0.2281±0.0140 |
| **Shuffle** | 0.6857±0.0077 | 0.3611±0.0052 | 0.0320±0.0026 |
| **- Volumes & flows** | 0.9590±0.0014 | 0.8326±0.0072 | 0.1760±0.0072 |
| **- Inh & exh kinetics** | 0.9537±0.0023 | 0.8326±0.0064 | 0.1901±0.0064 |
| **- Pauses after inh** | 0.9626±0.0020 | 0.8539±0.0049 | 0.1595±0.0049 |
| **-Pauses after exh** | 0.9596±0.0018 | 0.8519±0.0059 | 0.2416±0.0110 |
| **+ Local SDs** | 0.9589±0.0024 | 0.8857±0.0048 | 0.3326±0.0159 |

**Supplementary Table 3: pair-wise comparison of parameter exclusion conditions (Fig. 2B)**

a) **Wake** (N = 10 for each condition). Recall one-way ANOVA: F = 206.3205, df = 5, *P* = 6.7975e-34. Precision one-way ANOVA: F = 24.3523, df = 5, *P* = 9.6763e-13. Pairwise Dunnett test against 'All measurements' conditions:

| ***P*-values** | **Recall** | **Precision** |
|---|---|---|
| **- Volumes & flows** | 6.33561126e-04 | 4.78863045e-02 |
| **- Inh & exh kinetics** | 0.00000000e+00 | 1.76972881e-11 |
| **- Pauses after inh** | 8.49431636e-13 | 4.14999828e-01 |
| **-Pauses after exh** | 9.78989671e-01 | 1.87115931e-01 |
| **+ Local SDs** | 4.56808347e-09 | 3.63653170e-02 |

b) **NREM** (N = 10 for each condition). Recall one-way ANOVA: F = 326.2698, df = 5, *P* = 4.8575e-39. Precision one-way ANOVA: F = 117.9298, df = 5, *P* = 8.6897e-28. Pairwise Dunnett test against 'All measurements' conditions:

| ***P*-values** | **Recall** | **Precision** |
|---|---|---|
| **- Volumes & flows** | 0.00000000e+00 | 2.56852959e-07 |
| **- Inh & exh kinetics** | 3.03002168e-03 | 3.98989085e-07 |
| **- Pauses after inh** | 5.68166352e-07 | 1.41463975e-01 |
| **-Pauses after exh** | 1.40237268e-01 | 5.37968813e-01 |
| **+ Local SDs** | 0.00000000e+00 | 0.00000000e+00 |

c) **REM** (N = 10 for each condition). Recall one-way ANOVA: F = 75.8294, df = 5, *P* = 3.5884e-23. Precision one-way ANOVA: F = 316.8637, df = 5, *P* = 1.0416e-38. Pairwise Dunnett test against 'All measurements' conditions:

| **P-values** | **Recall** | **Precision** |
|---|---|---|
| **- Volumes & flows** | 9.99974791e-01 | 1.11022302e-15 |

| | | |
|---|---|---|
| **- Inh & exh kinetics** | 9.87861060e-01 | 1.77691772e-09 |
| **- Pauses after inh** | 0.00000000e+00 | 0.00000000e+00 |
| **-Pauses after exh** | 5.14461439e-01 | 3.64071132e-02 |
| **+ Local SDs** | 2.32344088e-04 | 0.00000000e+00 |

### Supplementary Table 4: Number of measurements for state transitions in Fig. 4D-F

| Transition | Amplitude PC1 | Sigmoid parameters PC1 | Amplitude PC3 | Sigmoid parameters PC3 |
|---|---|---|---|---|
| Wake to NREM | 672 | 343 | 672 | 0 |
| NREM to Wake | 524 | 340 | 524 | 211 |
| NREM to REM | 232 | 149 | 232 | 192 |
| REM to NREM | 92 | 64 | 92 | 71 |
| REM to Wake | 141 | 75 | 141 | 74 |

### Supplementary Table 5: Mean ± SD of PC1 at state transition for Fig. 4D-F

| Transition | Amplitude PC1 | Transition duration (cycle) | Delay to 10% change ($d_{onset}$) | Delay to 90% change ($d_{offset}$) |
|---|---|---|---|---|
| Wake to NREM | 0.7480 ± 0.5064 | 74.3260 ± 97.1537 | -54.5257 ± 69.6105 | 19.8003 ± 78.0731 |
| NREM to Wake | -0.8774 ± 0.5885 | 48.7972 ± 73.5383 | -6.2138 ± 49.0523 | 42.5833 ± 59.6425 |
| NREM to REM | 0.2378 ± 0.3333 | 49.6519 ± 61.8386 | -69.7223 ± 68.1611 | -20.0704 ± 60.7863 |
| REM to NREM | -0.4154 ± 0.4552 | 35.4366 ± 70.1857 | -12.6196 ± 61.5179 | 22.8170 ± 95.9832 |
| REM to Wake | -0.9792 ± 0.4961 | 38.8717 ± 60.0611 | 1.9040 ± 71.3610 | 40.7757 ± 77.2992 |

### Supplementary Table 6: Mean ± SD of PC3 at state transition for Fig. 4D-F

| Transition | Amplitude PC3 | Transition duration (cycle) | Delay to 10% change ($d_{onset}$) (cycle) | Delay to 90% change ($d_{offset}$) (cycle) |
|---|---|---|---|---|
| Wake to NREM | -0.2211 ± 0.5567 | not applicable | not applicable | not applicable |
| NREM to Wake | 0.4382 ± 0.7435 | 21.2508 ± 37.5871 | -9.9020 ± 58.1281 | 11.3488 ± 59.6098 |
| NREM to REM | -1.2124 ± 0.7709 | 91.2606 ± 79.4494 | -96.4055 ± 85.1001 | -5.1449 ± 63.6420 |
| REM to NREM | 1.3351 ± 0.8983 | 51.2010 ± 103.9029 | -20.4322 ± 53.5661 | 30.7688 ± 94.7925 |
| REM to Wake | 2.1273 ± 0.7607 | 46.3977 ± 104.7381 | -16.6520 ± 47.9260 | 29.7456 ± 64.0517 |

**Supplementary Table 7: ANOVA F, degree-of-freedom (df) and *P* values of transition parameters (Fig. 4D-F)**

| State transition | Amplitude | Transition duration | Delay to 10% change ($d_{onset}$) | Delay to 90% change ($d_{offset}$) |
|---|---|---|---|---|
| **PC1** | F=900.4, *P*=0.0000 (BRE) df=4 | F=3.06 *P*=0.0167 df=4 | F=20.31, *P*=2.27066e-15 df=4 | F=9.85, *P*=1.23476 e-7 df=4 |
| **PC3** | F=654.34, P=0.0000 df=4 | F=15.8, P=2.1936 3e-9 df=3 | F=27.82, *P*=2.05625 e-15 df=3 | F=2.04, *P*=0.1085 df=3 |

**Supplementary Table 8: Pairwise comparisons of amplitudes at state transitions with Tukey's Honest Significance Difference (HSD) tests (Fig. 4D)**

**PC1**

| Transition | NREM to Wake | NREM to REM | REM to NREM | REM to Wake |
|---|---|---|---|---|
| **Wake to NREM** | 0.0000 | 0.0000 | 0.0000 | 0.0000 |
| **NREM to Wake** | | 0.0000 | 0.0000 | 0.2194 |
| **NREM to REM** | | | 0.0000 | 0.0000 |
| **REM to NREM** | | | | 0.0000 |

**PC3**

| Transition | NREM to Wake | NREM to REM | REM to NREM | REM to Wake |
|---|---|---|---|---|
| **Wake to NREM** | 0.0000 | 0.0000 | 0.0000 | 0.0000 |
| **NREM to Wake** | | 0.0000 | 0.0000 | 0.0000 |
| **NREM to REM** | | | 0.0000 | 0.0000 |
| **REM to NREM** | | | | 0.0000 |

**Supplementary Table 9: Pairwise comparisons of state transition durations with HSD tests (Fig. 4E)**

**PC1**

| Transition | NREM to Wake | NREM to REM | REM to NREM | REM to Wake |
|---|---|---|---|---|
| **Wake to NREM** | 0.0300 | 0.2652 | 0.2625 | 0.4634 |
| **NREM to Wake** | | 1.0000 | 0.9593 | 0.9907 |
| **NREM to REM** | | | 0.9632 | 0.9904 |
| **REM to NREM** | | | | 0.9999 |

**PC3**

| Transition | NREM to REM | REM to NREM | REM to Wake |
|---|---|---|---|
| **NREM to Wake** | 0.000 | 0.1917 | 0.5468 |
| **NREM to REM** | | 0.0434 | 0.0888 |
| **REM to NREM** | | | 0.9964 |

**Supplementary Table 10: Pairwise comparisons of adaptation delay parameters with HSD tests (Fig. 4F)**

**PC1 $d_{onset}$**

| Transition | NREM to Wake | NREM to REM | REM to NREM | REM to Wake |
|---|---|---|---|---|
| Wake to NREM | 0.0000 | 0.4753 | 0.0337 | 0.0044 |
| NREM to Wake | | 0.0000 | 0.9924 | 0.9873 |
| NREM to REM | | | 0.0034 | 0.0004 |
| REM to NREM | | | | 0.9570 |

**PC1 $d_{offset}$**

| Transition | NREM to Wake | NREM to REM | REM to NREM | REM to Wake |
|---|---|---|---|---|
| Wake to NREM | 0.0167 | 0.0011 | 0.8878 | 0.7699 |
| NREM to Wake | | 0.0000 | 0.3229 | 1.0000 |
| NREM to REM | | | 0.9481 | 0.0149 |
| REM to NREM | | | | 0.5702 |

**PC3 $d_{onset}$**

| Transition | NREM to REM | REM to NREM | REM to Wake |
|---|---|---|---|
| NREM to Wake | 0.000 | 0.8897 | 0.9833 |
| NREM to REM | | 0.0000 | 0.0001 |
| REM to NREM | | | 0.9981 |

**PC3 $d_{offset}$**

| Transition | NREM to REM | REM to NREM | REM to Wake |
|---|---|---|---|
| NREM to Wake | 0.2458 | 0.9635 | 0.6812 |
| NREM to REM | | 0.9495 | 0.1558 |
| REM to NREM | | | 0.6141 |

**Supplementary Table 11: Summary table of mice/sessions used for statistical comparisons in Fig. 1.**

Overall N =38 sessions from N = 8 mice were used to compare respiration between states: mouse 3C060 = 6 sessions, mouse 3C209 (female) = 1 session, mouse 3C028 = 3 sessions, mouse 7C026 = 2 sessions, mouse 7C012 = 4 sessions, mouse 7C014 = 4 sessions, mouse 3C030 = 14 sessions, mouse 3C083 = 3 sessions.

The overall duration in seconds and the corresponding number of cycles are reported for each state (Wake, NREM, REM).

| | | Duration (seconds) | | | N cycles | | |
|---|---|---|---|---|---|---|---|
| **Mouse** | **Session** | **Wake** | **NREM** | **REM** | **Wake** | **NREM** | **REM** |
| 3C060 | 3C060-S11 | 1490 | 1422 | 177 | 7742 | 4673 | 650 |
| 3C060 | 3C060-S12 | 3992 | 7766 | 1093 | 22536 | 23724 | 3942 |
| 3C060 | 3C060-S16 | 8471 | 6706 | 771 | 45428 | 19163 | 2722 |
| 3C060 | 3C060-S17 | 9290 | 7548 | 570 | 50300 | 22247 | 2014 |
| 3C060 | 3C060-S18 | 10325 | 7094 | 879 | 55391 | 18987 | 3084 |
| 3C060 | 3C060-S19 | 9313 | 8049 | 727 | 51852 | 22882 | 2645 |
| 3C209 | 3C209_S16 | 5986 | 14290 | 1196 | 32284 | 36933 | 4193 |
| 3C028 | 3C028_S28 | 9213 | 3939 | 427 | 61546 | 11237 | 1393 |
| 3C028 | 3C028_S30 | 8821 | 6392 | 356 | 56928 | 18442 | 1243 |
| 3C028 | 3C028_S31 | 9418 | 4337 | 324 | 60903 | 13998 | 1147 |
| 7C026 | 7C026-S27 | 9502 | 8432 | 989 | 46352 | 17027 | 2992 |
| 7C026 | 7C026-S31 | 5512 | 10099 | 1133 | 23922 | 22177 | 3631 |
| 7C012 | 7C012-S11 | 7576 | 8284 | 838 | 33841 | 22295 | 2675 |
| 7C012 | 7C012-S12 | 5396 | 8406 | 524 | 22408 | 20931 | 1618 |
| 7C012 | 7C012-S16 | 8462 | 8014 | 670 | 42009 | 24664 | 2236 |
| 7C012 | 7C012-S22 | 7503 | 5917 | 571 | 30531 | 15358 | 1995 |
| 7C012 | 7C012-S27 | 9669 | 6498 | 630 | 36677 | 20100 | 2198 |
| 7C014 | 7C014-S10 | 6585 | 9586 | 679 | 27040 | 27543 | 2358 |
| 7C014 | 7C014-S11 | 7428 | 9400 | 404 | 30049 | 25524 | 1493 |
| 7C014 | 7C014-S12 | 8225 | 7647 | 360 | 35318 | 21846 | 1228 |
| 7C014 | 7C014-S17 | 8229 | 3055 | 51 | 37540 | 5114 | 118 |
| 3C030 | 3C030-S10_191126 | 10234 | 3786 | 310 | 62349 | 7517 | 833 |
| 3C030 | 3C030-S11_191127 | 11251 | 5747 | 327 | 60400 | 10550 | 807 |
| 3C030 | 3C030-S12_191128 | 12912 | 6700 | 539 | 71588 | 12112 | 1434 |
| 3C030 | 3C030-S18_191208 | 13419 | 4449 | 252 | 76549 | 9066 | 671 |
| 3C030 | 3C030-S19_191209 | 10023 | 6152 | 388 | 55975 | 12737 | 1137 |

| | | | | | | | |
|---|---|---|---|---|---|---|---|
| 3C030 | 3C030-S20_191210 | 12014 | 6444 | 495 | 66388 | 13443 | 1274 |
| 3C030 | 3C030-S22_191212 | 10102 | 6625 | 499 | 51780 | 15108 | 1414 |
| 3C030 | 3C030-S23_191213 | 11919 | 5613 | 173 | 55853 | 9608 | 478 |
| 3C030 | 3C030-S24_191214 | 11411 | 4419 | 140 | 54829 | 7797 | 432 |
| 3C030 | 3C030-S25_191215 | 10604 | 5558 | 324 | 49928 | 9751 | 1040 |
| 3C030 | 3C030-S26_191216 | 9983 | 6613 | 243 | 45778 | 13367 | 707 |
| 3C030 | 3C030-S27_191217 | 10444 | 5438 | 234 | 51008 | 12032 | 680 |
| 3C030 | 3C030-S28_191218 | 11646 | 4958 | 271 | 57140 | 8252 | 781 |
| 3C030 | 3C030-S29_191219 | 11582 | 8671 | 463 | 52273 | 15802 | 1263 |
| 3C083 | 3C083_S22_200909 | 12034 | 7789 | 240 | 52186 | 12544 | 550 |
| 3C083 | 3C083_S33_200926 | 13026 | 4827 | 39 | 53191 | 6961 | 79 |
| 3C083 | 3C083_S35_200928 | 10461 | 6940 | 292 | 45971 | 10272 | 608 |

### Supplementary figures

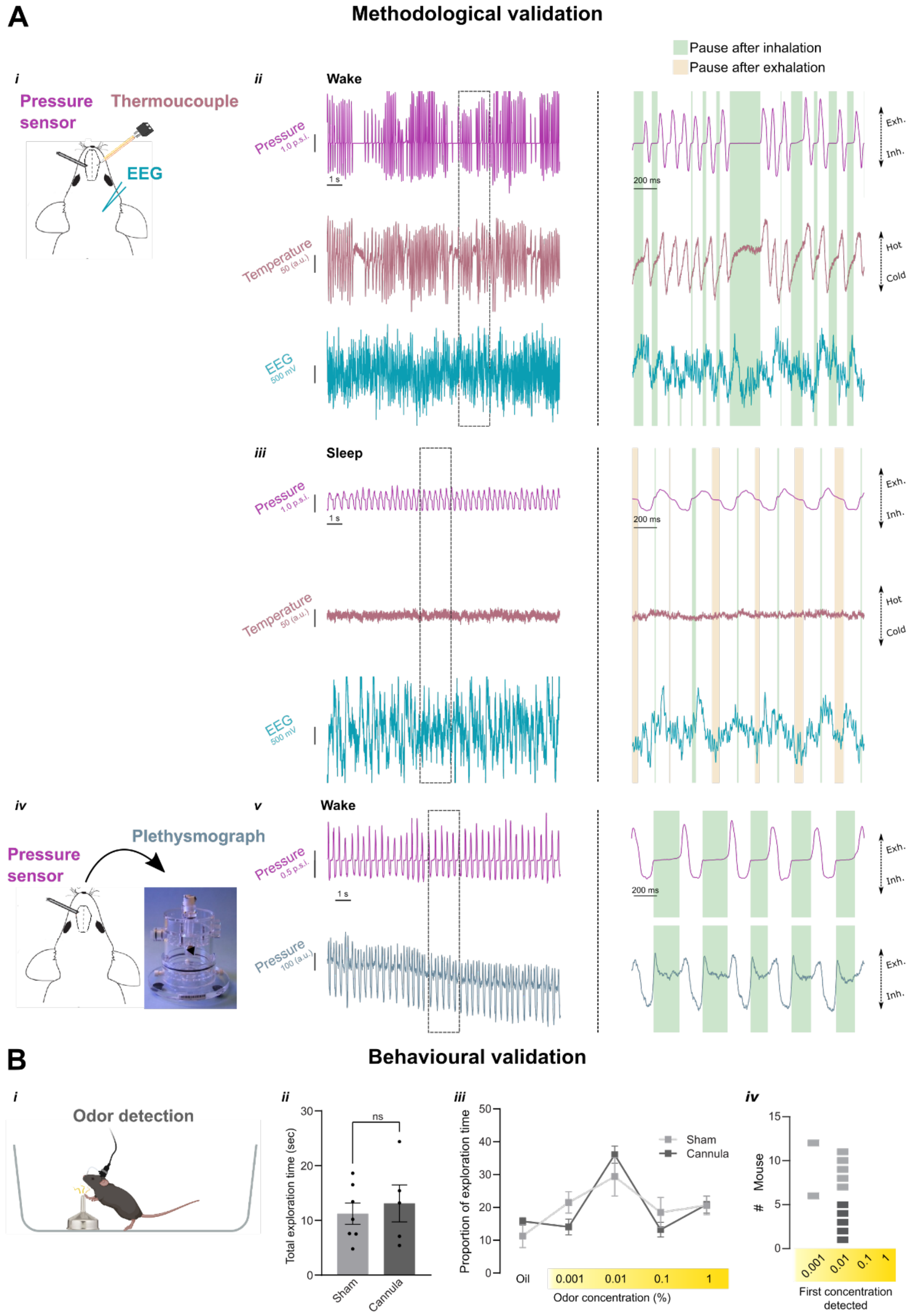

**Supplementary Figure 1: Validation of pressure signal recordings for monitoring respiration in freely-moving mice.**

**A**. Methodological validation comparing respiration signal recorded with pressure sensor with that of thermocouple and plethysmograph.

***i)*** Schematic of the experimental procedure. The nasal cannula connected to the pressure sensor is chronically implanted into the left nasal cavity while the thermocouple is implanted into the right one. A screw above the frontal cortex is used for EEG recording and one is placed above the cerebellum as reference for the EEG recordings. All signals are simultaneously acquired in a freely moving mouse.

***ii)*** Color-coded examples of raw recordings (left = 15 seconds, right = 2 seconds) collected in parallel during Wake showing variations in nasal pressure (*top*) or intranasal temperature (*middle*) in arbitrary units during simultaneous EEG recording (*bottom*). Note the overall similarity between respiration signals obtained with pressure sensor and thermocouple during both inhalation/exhalation and the lack of detection of pauses after inhalation (green area) with thermocouples (see zoomed in panel on the right).

***iii)*** Same as for Wake, color-coded examples of raw recordings during sleep. Note the attenuated, yet present, respiration signal obtained with pressure sensor in contrast to the overall flattening with the thermocouples. Same as for Wake, respiratory pauses can be detected only with pressure sensors unlike thermocouples (see zoomed in panel on the right).

***iv)*** Schematic of the experimental procedure. The nasal cannula connected to the pressure sensor is chronically implanted into the left nasal cavity. The mouse was placed in the chamber of a plethysmograph while nasal pressure was simultaneously monitored. ***v)*** Examples of raw recordings (left = 15 seconds, right = 2 seconds) collected in parallel during Wake state showing variations in nasal pressure (*top*) or plethysmograph pressure (*bottom*). Note the overall similarity between respiration signals obtained with pressure sensor and plethysmography during both inhalation/exhalation and the pauses after inhalation (see zoomed in panel on the right).

**B**. Odor detection task in mice implanted with nasal cannula. ***i)*** Illustration of the explorative odor detection task (see Methods). ***ii)*** Proportion of exploration time (i.e. spent sniffing mineral oil and odors) for sham (grey, N = 7) and cannula-implanted (Cannula, dark grey, N = 5) mice. Sham mean 11.23 ± 1.93 sec vs. Cannula mean 13.10 ± 3.38 sec (Mann-Whitney rank sum test: $P > 0.999$). ***iii)*** Mean ± SEM total exploration time within the 5 trials comparing Sham (grey, N =7) and Cannula (dark grey, N = 5 mice). Two-way repeated measures ANOVA interaction: $P = 0.395$. iv) Distribution of individual mice (Sham, grey, N = 7 and Cannula, dark grey, N = 5) according to the concentration they detected first.

**Supplementary Figure 2: Nasal pressure recordings from freely-moving mice across vigilance states**

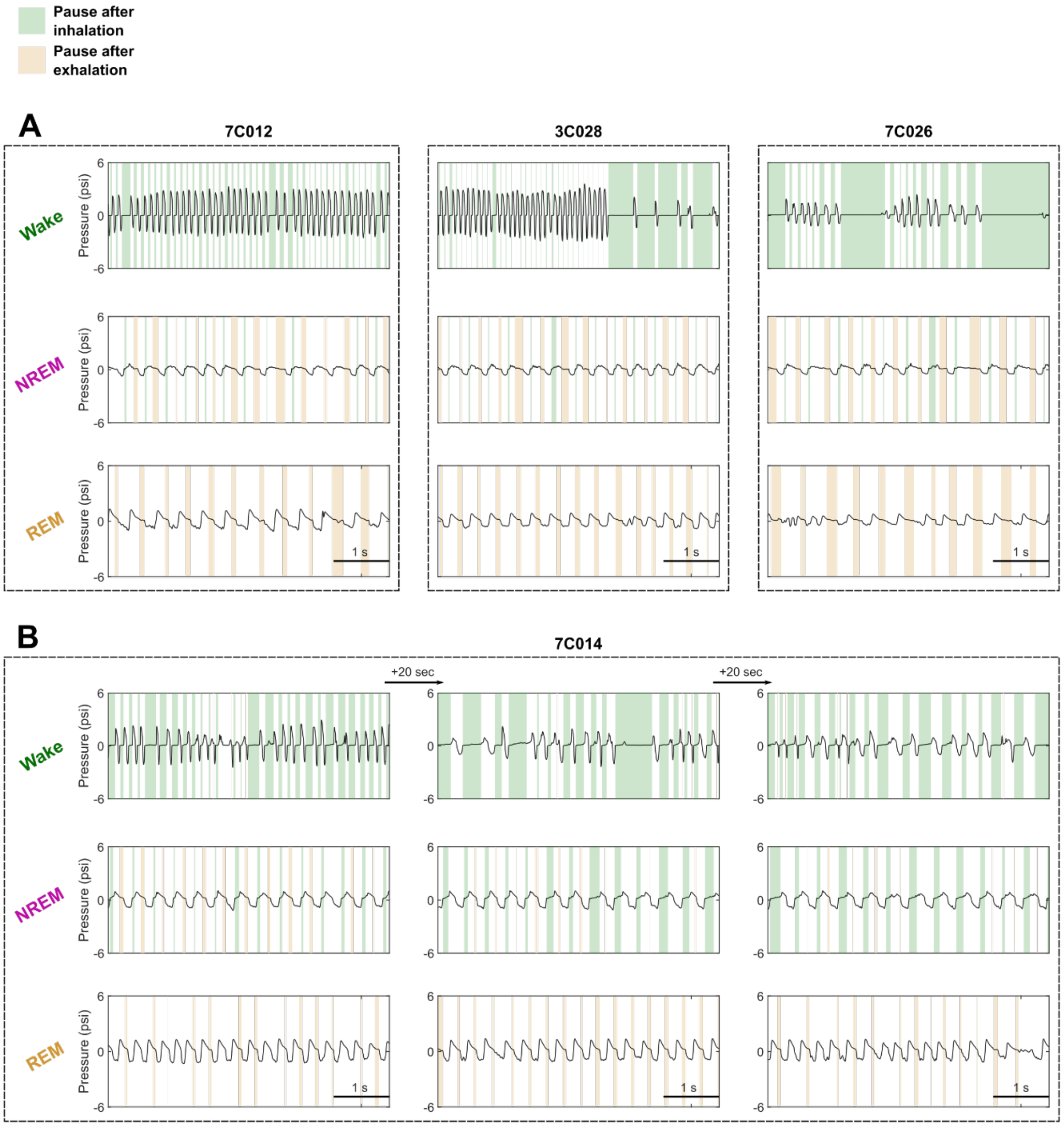

**A.** Representative examples of respiratory traces (5 seconds) from 3 mice (7C12, 3C028, 7C026) across Wake (*top*), NREM (*middle*) and REM states (*bottom*). The black line shows intranasal pressure signal, colour-coded areas represent pauses after inhalation (green) and pauses after exhalation (orange). Note the similarity in the pause respiratory patterns across mice while they overall differed across states: Wake and REM had the most profound differences in the location of pauses (Wake only pauses after inhalation, REM only pauses after exhalation), while NREM showed more intermediate patterns with both types of pauses.

**B.** Representative examples of respiratory traces as in **A** from mouse 7C014 showing respiratory pause patterns within a session across time (columns, time gap = 20 seconds). Note the stability in pause respiratory patterns during Wake and REM unlike NREM showing changes in the duration of pauses.

### Supplementary Figure 3: Pressure sensor analysis pipeline.

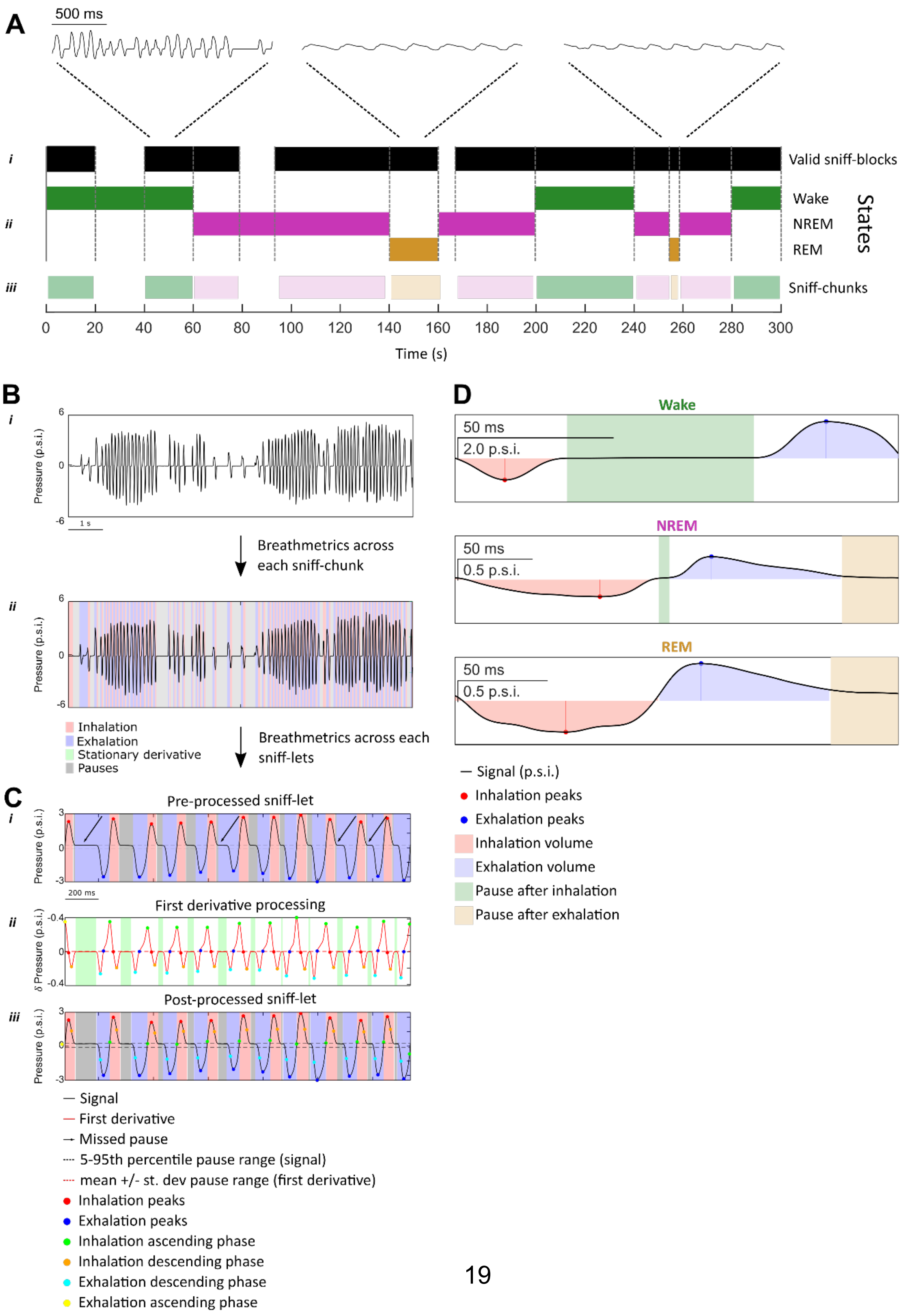

**A**. Schematic representation of a putative respiratory session recorded across time. ***i***) Valid sniff-blocks (black area) represent intervals during which respiration was considered as real. ***ii***) Color-coded intervals representing states (Wake = green, REM = orange, NREM = purple) were intercepted with valid sniff-blocks to produce sniff-chunks ***iii***) – consecutive intervals units of valid respiration assigned to an individual state.

**B**. Representative example of a single sniff-chunk assigned to wake state. ***i***) individually analysed with BreathMetrics. ***ii***) returning per-cycle information (inhalation, exhalation, pause intervals, peaks, through etc) across the whole sniff-chunk.

**C**. Schematic representation of the analytical pipeline used for each individual sniff-let. ***i***) Representative example of an individual sniff-let comprising 11 respiration cycles prior to re-iterative BreathMetrics analysis. ***ii***) First derivative of the respiration signal of the corresponding sniff-let shown in ***i***) highlight minima/maxima (representing inhalation ascending phase, inhalation descending phase, exhalation descending phase and exhalation ascending phase) alternated to stationary epochs (green area). ***iii***) Same sniff-let shown in (*i*) after re-iterative BreathMetrics analysis showed refined inhalation, exhalation and pause intervals.

**D.** Examples of respiration cycles in the three brain states. Different x-y(time-pressure) scales have been used to highlight the architecture of respiration cycles composed of inhalation, exhalation, intermingled with pauses. For both inhalation and exhalation, color-coded arrows represent time and amplitude of the peaks displayed as dots, while the areas represent the overall volume.

**Supplementary Figure 4: Comparisons of inhalation/exhalation features across brain states.**

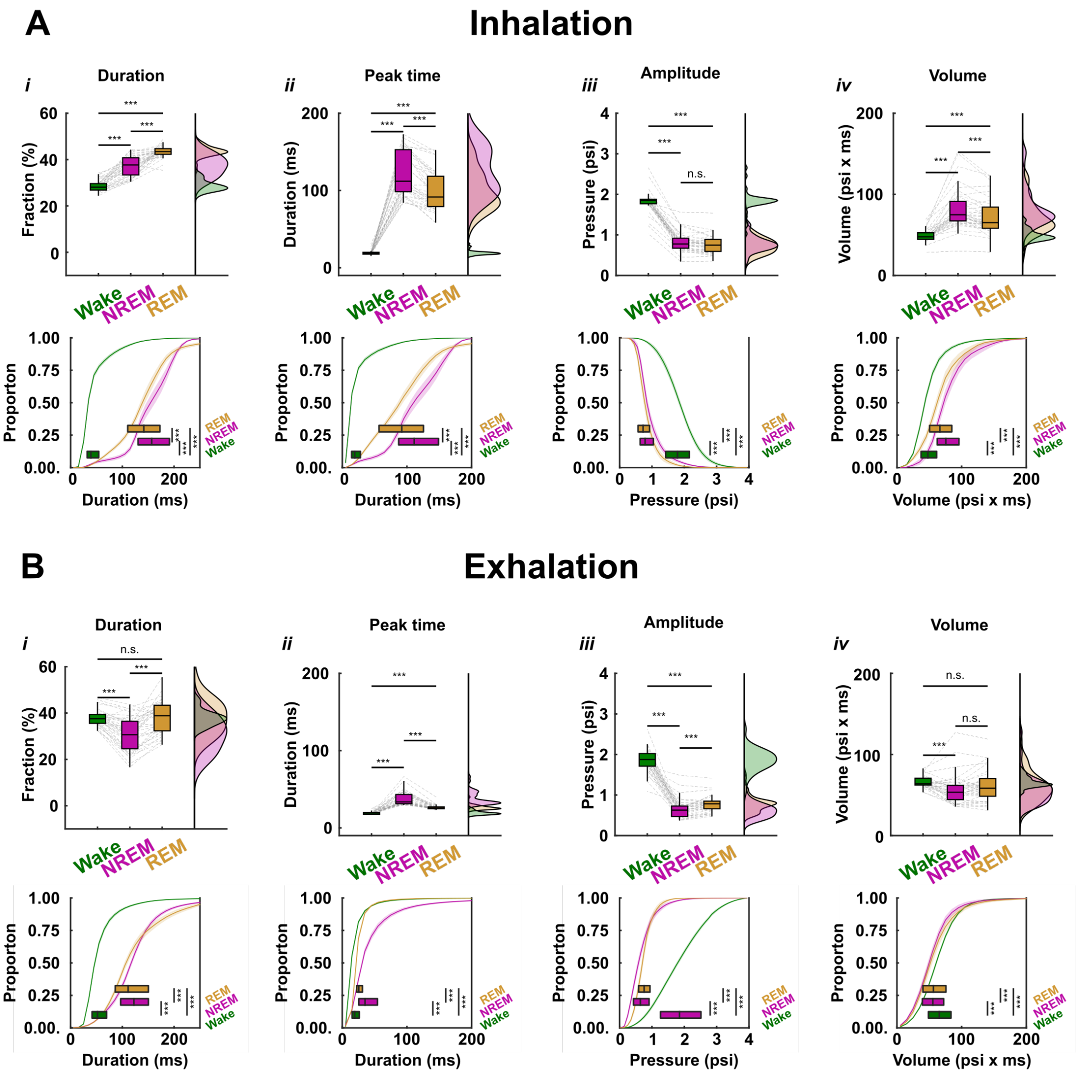

Descriptive statistics of the respiratory features (relative duration (%), peak time, amplitude and volume) for inhalation (**A**) and exhalation (**B**) during Wake (Green), NREM (purple) and REM (gold). Top rows show box plots representing distribution of the median values obtained across experimental sessions (*N* = 38) and compared with non-parametric test (Kruskal-Wallis test followed by post-hoc Wilcoxon test with Bonferroni correction alpha = 0.05/3). Bottom rows show both the averaged cumulative distributions, compared with two-sample Kolmogorov-Smirnov tests and overimposed boxplots obtained by pooling values from random 500 cycles across all sessions and then compared with non-parametric test (Kruskal-Wallis test followed by *post-hoc* Wilcoxon test with Bonferroni correction alpha = 0.05/3).

**A.** Respiratory features pertaining to inhalation in Wake, NREM and REM.

***i)*** Relative inhalation duration (% of the total cycle duration):
*(Top)* Comparisons of the median inhalation percentage across sessions revealed a significant decrease during Wake as compared to both REM and NREM and a significant decrease during NREM compared to REM. Inhalation time ± SEM: wake = 26.7 ± 0.4 %, NREM = 37.3 ± 0.7 %, REM = 43.5 ± 0.4 %, Kruskal-Wallis test: $\chi^2(2) = 87.1$, $P = 1.23e\text{-}19$, *post-hoc* Wilcoxon test, all $P < 0.0001$.
(*Bottom*) Two samples Kolmogorov-Smirnov test: Wake vs NREM: D(36214) = 0.39, $P < 0.0001$; Wake vs REM: D(36214) = 0.63, $P < 0.0001$, NREM vs REM: D(36214) = 0.39, $P < 0.0001$; Kruskal-Wallis test: $\chi^2(2) = 18107$, $P = 2.22e\text{-}16$, *post-hoc* Wilcoxon test, all: $P < 0.00001$.
***ii)*** Inhalation peak time:
*(Top)* Comparisons of the median inhalation peak time revealed a significant decrease during Wake as compared to both NREM and NREM and a significant increase in NREM compared to REM. Inhalation peak time ± SEM: wake = 19.5 ± 0.4 ms, NREM = 123.0 ± 0.5 ms, REM = 98.4 ± 4.0 ms, Kruskal-Wallis test: $\chi^2(2) = 81.7$, $P = 1.83e\text{-}18$, *post-hoc* Wilcoxon test with Bonferroni correction, all $P < 0.0001$.
(*Bottom*) Two samples Kolmogorov-Smirnov test: Wake vs NREM: D(36214) = 0.81, $P = 1.22e\text{-}16$; Wake vs REM: D(36214) = 0.71, $P = 1.32e\text{-}16$, NREM vs REM: D(36214) = 0.23, $P = 4.12e\text{-}16$; Kruskal-Wallis test: $\chi^2(2) = 28378$, $P = 6.65e\text{-}16$, *post-hoc* Wilcoxon test, all: $P < 0.00001$.
***iii)*** Inhalation amplitude:
*(Top)* Comparisons of the median inhalation amplitude revealed a significant increase during Wake compared to both NREM and REM and no differences during NREM

compared to REM. Inhalation peak amplitude mean ± SEM: Wake = -1.82 ± 0.04 p.s.i., NREM = -0.81 ± 0.04 p.s.i., REM -0.76 ± 0.04 p.s.i., Kruskal-Wallis test: $\chi^2(2) = 74.0$, $P$ = 8.64e-17, *post-hoc* Wilcoxon test with Bonferroni correction, Wake vs NREM: $P$ =7.73e-08, Wake vs REM: $P$ = 7.73e-08, NREM vs REM: $P$ = 0.021.

(*Bottom*) Two samples Kolmogorov-Smirnov test: Wake vs NREM: D(36214) = 0.69, $P$ = *1*.42e-15; Wake vs REM: D(36214) = 0.76, $P$ = 5.65e-15, NREM vs REM: D(36214) =2.21 0.11, $P$ = 2.21e-18; Kruskal-Wallis test: $\chi^2(2) = 26337$, $P$ = 6.90e-16, *post-hoc* Wilcoxon test, all: $P < 0.00001$.

***iv)*** Inhalation volume:

*(Top*) Comparisons of the median inhalation volume revealed a significant decrease in Wake compared to NREM and REM and a significant increase in NREM compared to REM. Inhalation volume ± SEM: wake = 48.8 ± 1.3 p.s.i. x ms, NREM = 81.6 ± 4.0 p.s.i. x ms, REM = 72.3 ± 3.5 p.s.i. x ms, Kruskal-Wallis test: $\chi^2(2) = 56.9$, $P$ = 4.33e-13, *post-hoc* Wilcoxon test with Bonferroni correction, all $P < 0.0001$.

(*Bottom*) Two samples Kolmogorov-Smirnov test: Wake vs NREM: D(36214) = 0.52, $P$ = 1.56e-16; Wake vs REM: D(36214) = 0.34, $P$ = 5.71e-16, NREM vs REM: D(36214) = 0.19, $P$ =1.76e-16; Kruskal-Wallis test: $\chi^2(2) = 10721$, $P$ = 9.16e-15, *post-hoc* Wilcoxon test, all: $P < 0.0001$.

**B**. Respiratory features pertaining to inhalation in Wake, NREM and REM.

***i)*** Relative exhalation duration (% of the total cycle duration):

*(Top)* Comparisons of the median exhalation percentage across sessions revealed a significant decrease during Wake as compared to NREM but not REM and a significant decrease during NREM compared to REM. Mean exhalation time ± SEM: wake = 37.9 ± 0.6 %, NREM = 30.9 ± 1.2 %, REM = 38.1 ± 1.1 %, Kruskal-Wallis test: $\chi^2(2) = 22.0$, $P$ = 1.65e-05, *post-hoc* Wilcoxon test, Wake vs NREM: $P$ = 3.44e-07, Wake vs REM: $P$ = 0.89, NREM vs REM: $P$ = 7.32e-07.

(*Bottom*) Two samples Kolmogorov-Smirnov test: Wake vs NREM: D(36214) = 0.18, $P$ = 1.61e-16; Wake vs REM: D(36214) = 0.11, $P$ = 7.84e-16, NREM vs REM: D(36214) = 0.18, $P$ = 2.79e-15; Kruskal-Wallis test: $\chi^2(2) = 2446$, P = 2.35e-16, post-hoc Wilcoxon test, all: $P < 0.00001$.

***ii)*** Exhalation peak time:

*(Top)* Comparisons of the median exhalation peak time revealed a significant decrease during Wake as compared to both NREM and NREM and a significant increase in NREM compared to REM. Exhalation peak time ± SEM: wake = 19.5 ± 0.4 ms, NREM = 38.0 ± 1.4 ms, REM = 26.4 ± 0.5 ms, Kruskal-Wallis test: $\chi^2(2) = 94.9$, *P = 2.43e-21*, *post-hoc* Wilcoxon test with Bonferroni correction, all $P < 0.0001$.

(*Bottom*) Two samples Kolmogorov-Smirnov test: Wake vs NREM: D(36214) = 0.48, *P* = 9.81e-16; Wake vs REM: D(36214) = 0.40, *P =1.*22e-16, NREM vs REM: D(36214) = 0.34, *P* = 5.01e-16; Kruskal-Wallis test: $\chi^2(2) = 11136$, *P* = 1.36e-17, *post-hoc* Wilcoxon test, all: $P < 0.0001$.

***iii)*** Exhalation amplitude:

*(Top)* Comparisons of the median exhalation amplitude revealed a significant increase during Wake compared to both NREM and REM and a significant decrease during NREM compared to REM. Exhalation peak amplitude mean ± SEM: Wake = 1.86 ± 0.04 p.s.i., NREM = 0.65 ± 0.04 p.s.i., REM = 0.78 ± 0.03 p.s.i., Kruskal-Wallis test: $\chi^2(2) = 79.1$, *P* = 6.54e-18, *post-hoc* Wilcoxon test with Bonferroni correction, all $P < 0.0001$.

(*Bottom*) Two samples Kolmogorov-Smirnov test: Wake vs NREM: D(36214) = 0.68, *P* = 6.74e-16; Wake vs REM: D(36214) = 0.69, *P* = 6.89e-16, NREM vs REM: D(36214) = 0.20, *P* = 3.35e-16; Kruskal-Wallis test: $\chi^2(2) = 24073$, *P* = 2.51e-16, *post-hoc* Wilcoxon test, all: $P < 0.0001$.

***iv)*** Exhalation volume:

*(Top*) Comparisons of the median exhalation volume revealed a significant increase in Wake compared to NREM but not REM and no significant changes between NREM and REM. Exhalation volume ± SEM: wake = 65.9 ± 1.0 p.s.i. x ms, NREM = 56.7 ± 2.9 p.s.i. x ms, REM = 60.6 ± 3.1 p.s.i. x ms, Kruskal-Wallis test: $\chi^2(2) = 16.5$, *P* =0.0003, *post-hoc* Wilcoxon test with Bonferroni correction, Wake vs NREM: *P* = 0.0003, Wake vs REM: *P* = 0.021, NREM vs REM: *P* = 0.12.

(*Bottom*) Two samples Kolmogorov-Smirnov test: Wake vs NREM: D(36214) = 0.16, *P* = 8.08e-15; Wake vs REM: D(36214) = 0.15, *P* = 9.95e-16, NREM vs REM: D(36214) = 0.05, *P* = 2.44e-16; Kruskal-Wallis test: $\chi^2(2) = 1324$, *P* = 6.43e-16, *post-hoc* Wilcoxon test, all: $P < 0.0001$.

### Supplementary Figure 5: Distribution of input respiratory features used for brain state prediction.

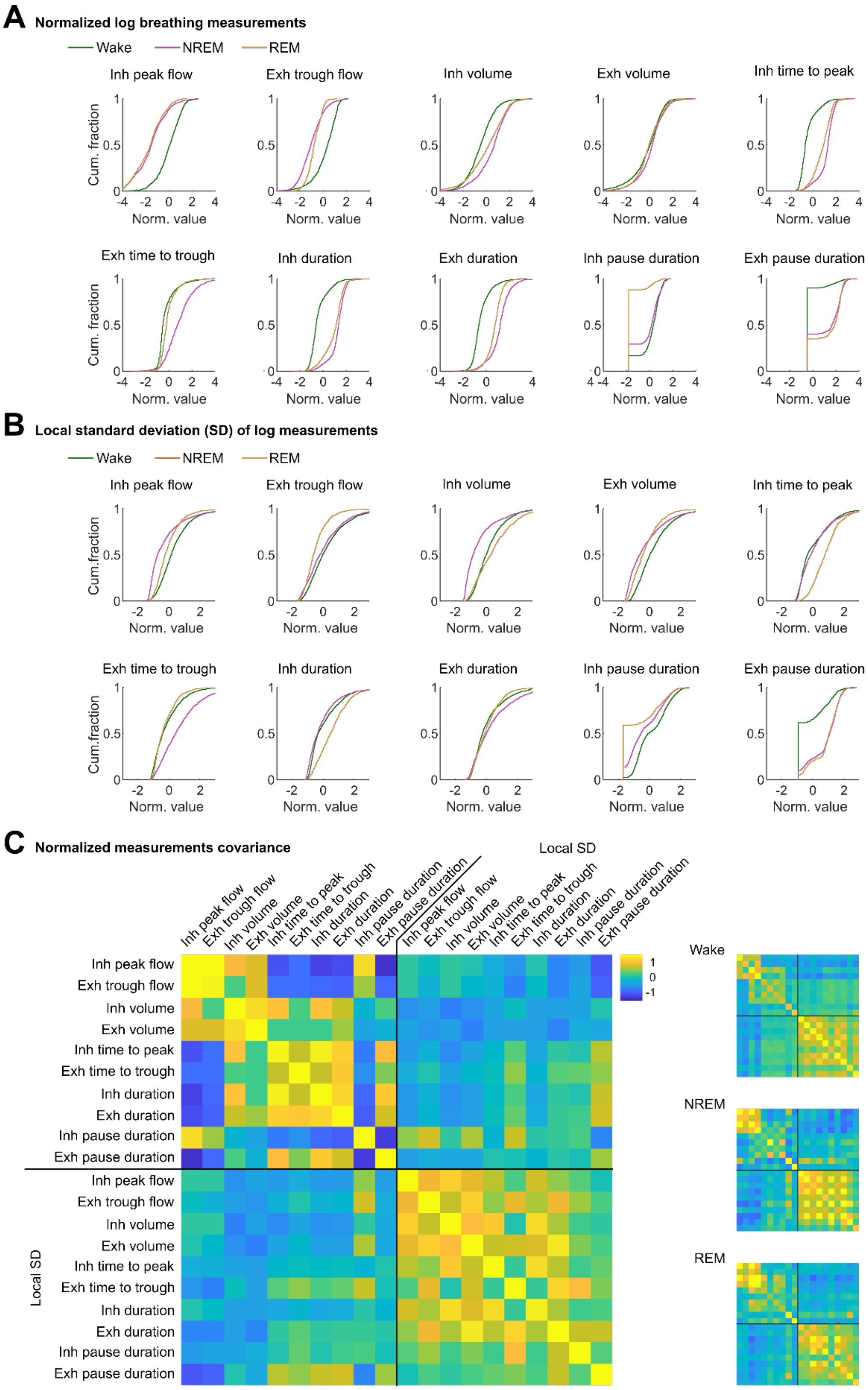

**A.** Cumulative distributions of log- and unit variance-normalized features (**Methods**) of the respiratory cycle (2000 sampled cycles for each of the 7 animals and each brain state). Inh: inhalation. Exh: exhalation.

**B.** For the same respiratory cycle as in **A**, distributions of the local standard deviation (SD) (over a window of width 10 cycles) of the normalized features.

**C.** Covariance matrix of all normalized features and their local standard deviation with equi-probable brain states (left) and for each brain state separately (right).

**Supplementary Figure 6: Artificial neuronal network for brain state prediction from respiratory features.**

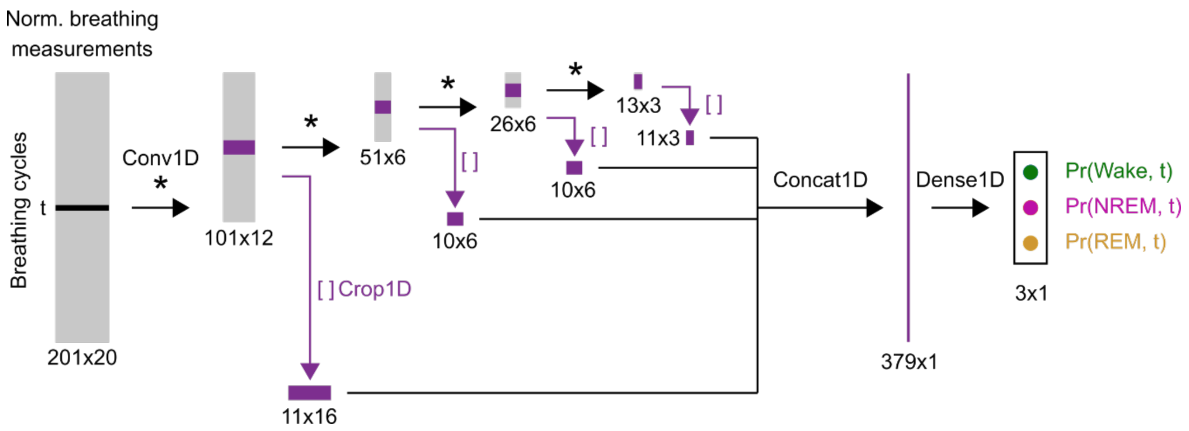

**A.** Structure of the artificial neuronal network used for state prediction in **Fig. 2**. The 20 input features (normalized features of the respiratory cycle and their local standard deviation, **Supplementary Table 1**) are taken over a 201 cycles time window around the cycle of interest (time *t*) and concatenated in a tensor of size 201x20. The network output is the probability of each of the three brain states for the respiratory cycle *t*. Conv1D (*): 1D convolutional layer. Crop1D ([]): central cropping operation. Concat1D: concatenate all coefficients in a 1D tensor. Dense1D: densely connected layer in 1D. All Conv1D layers used a bias and a ReLu activation function. The Dense1D layer used a bias and used a SoftMax activation function.

**Supplementary Figure 7: Principal component analysis of log-normalized features from respiratory cycles.**

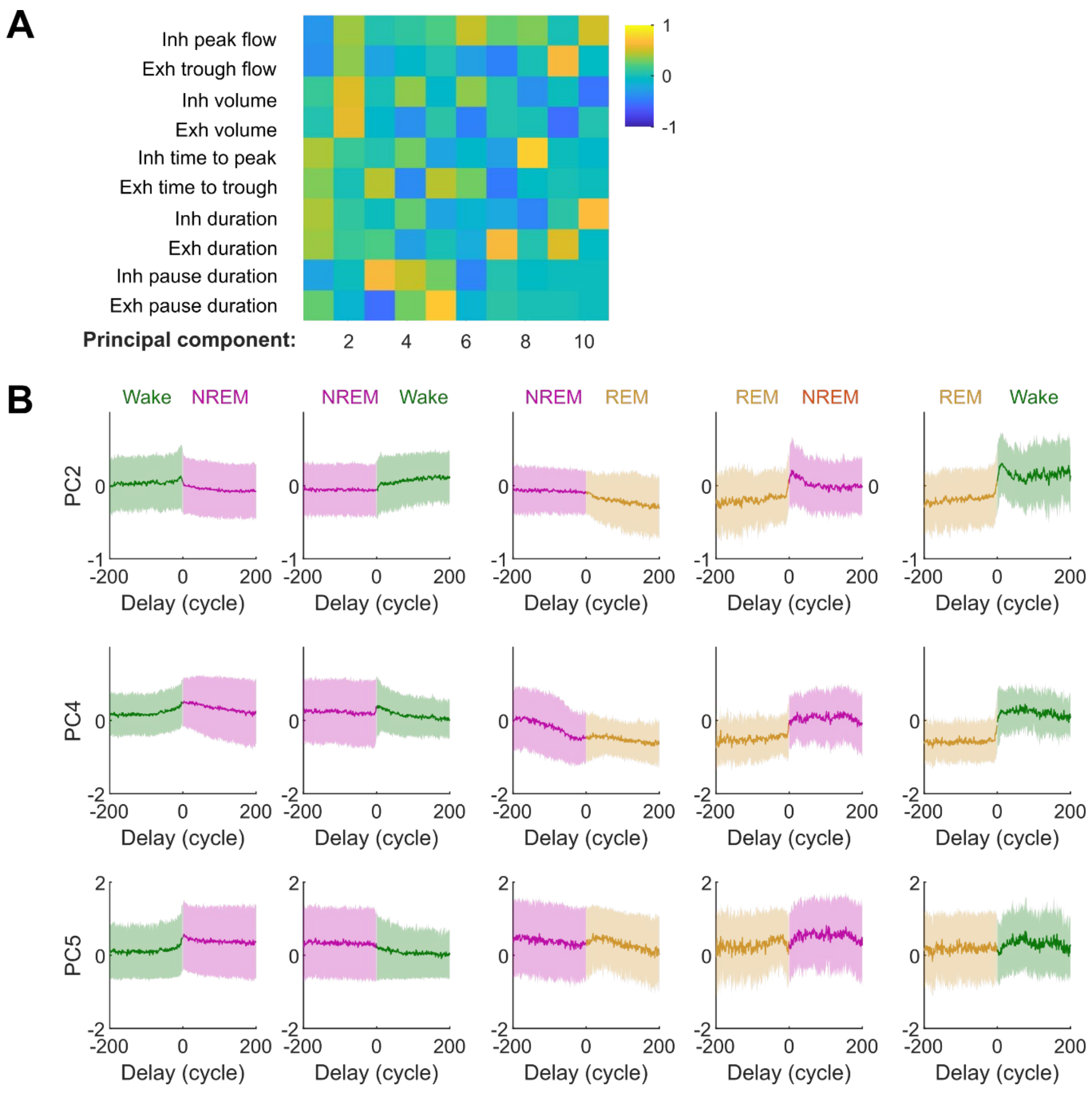

**A.** All components computed from PCA of the 10 features of respiration cycle features (after log- and unit variance normalization of individual features). Inh: inhalation. Exh: exhalation. Principal components are sorted, from left to right, in order of decreasing associated eigenvalue. Sample respiratory cycles were balanced in between brain states

(Wake, NREM, REM) and the 7 animals (2000 random cycles for each state-animal pair). **B.** Population averages ± standard deviation of coefficients for principal components #2, 4 and 5 at brain state transitions. Same events as for **Fig. 4G, H.** Wake→NREM N = 672; NREM→Wake N = 524; NREM→REM N = 232; REM→NREM N = 92, REM→Wake N = 141.

**Supplementary Figure 8: Dynamics of cycle-based respiratory features at cortical state transitions**

Population averages ± standard deviation of respiratory features (without normalization) at brain state transitions. Same events as for **Fig. 4G, H** and **Supplementary Fig. 7**. Inh: inhalation. Exh: exhalation.

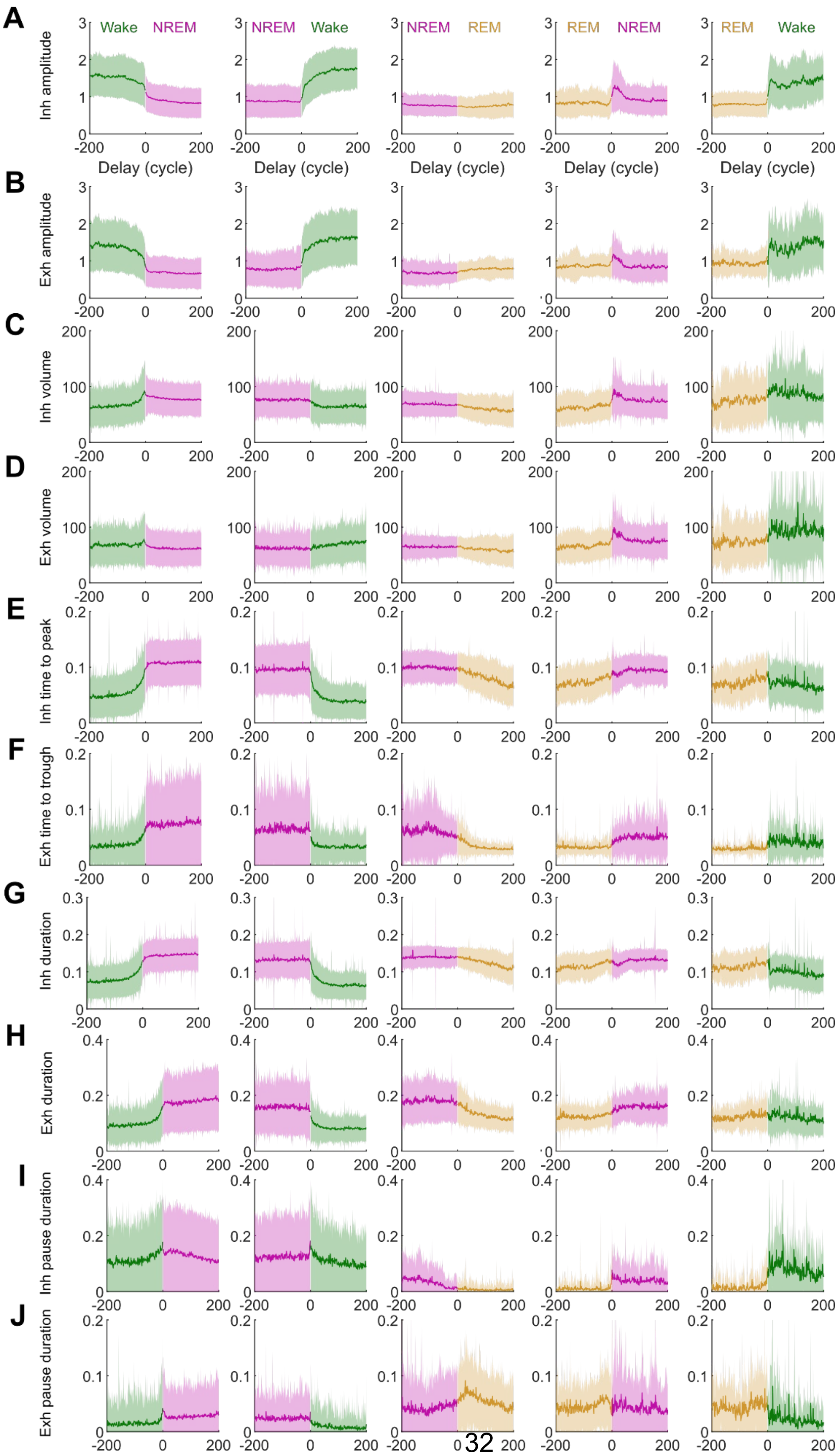

A
B
C
D
E
F
G
H
I
J
Wake
NREM
REM
Inh amplitude
Exh amplitude
Inh volume
Exh volume
Inh time to peak
Exh time to trough
Inh duration
Exh duration
Inh pause duration
Exh pause duration
Delay (cycle)

**Supplementary Figure 9: Dynamics of the local SD of cycle-based respiratory features at cortical state transitions**

**A-J.** Population averages ± standard deviation of the local (10 cycles) standard deviation (SD) of respiratory features (without normalization) at brain state transitions. Same events as for **Fig. 4D, E** and **Supplementary Fig. 7 and 8**. Inh: inhalation. Exh: exhalation.

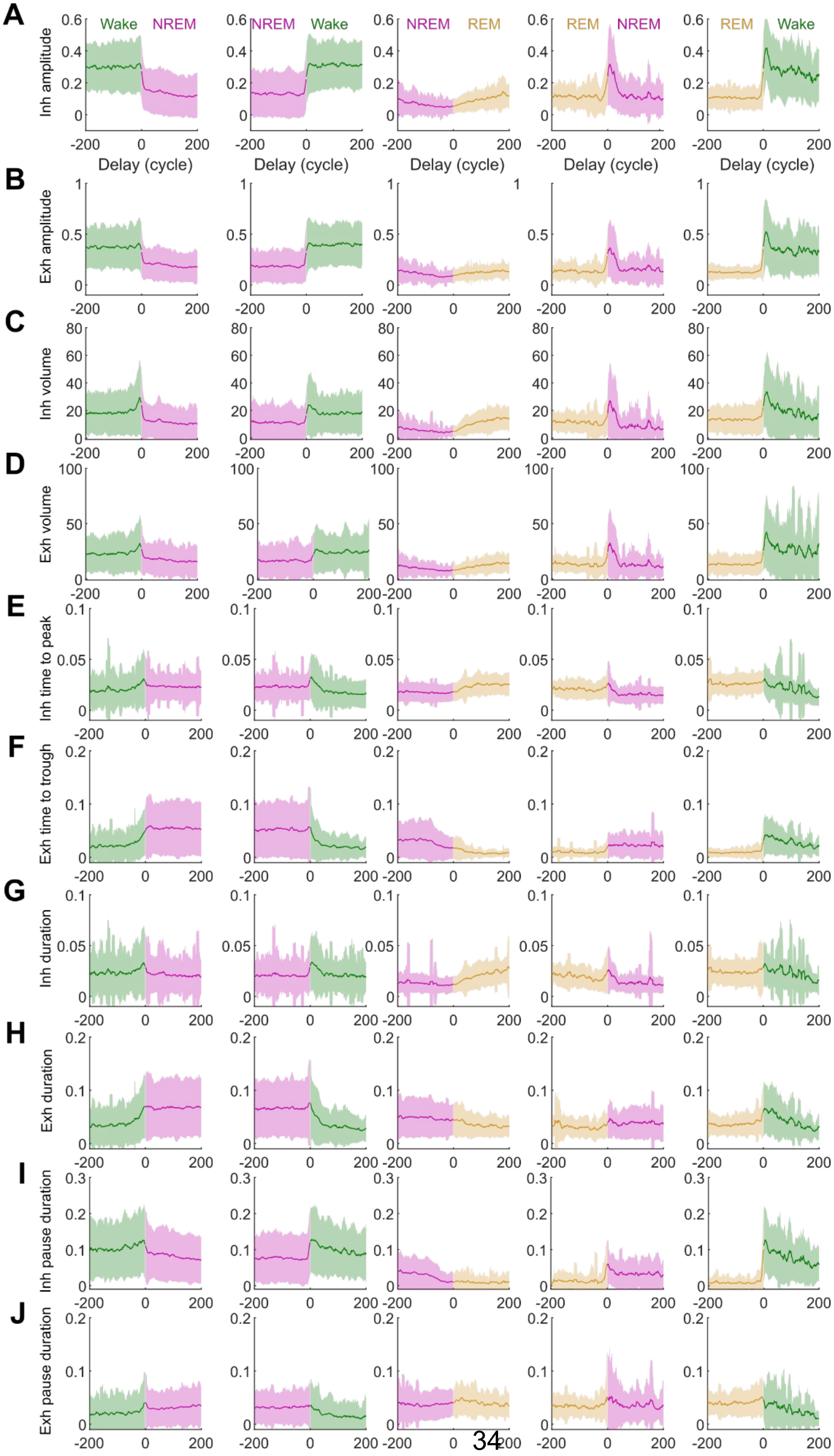

A
B
C
D
E
F
G
H
I
J
Wake
NREM
REM
Inh amplitude
Exh amplitude
Inh volume
Exh volume
Inh time to peak
Exh time to trough
Inh duration
Exh duration
Inh pause duration
Exh pause duration
Delay (cycle)

**Supplementary Figure 10: Respiratory features during normalized NREM packets and at transitions with microarousals.**

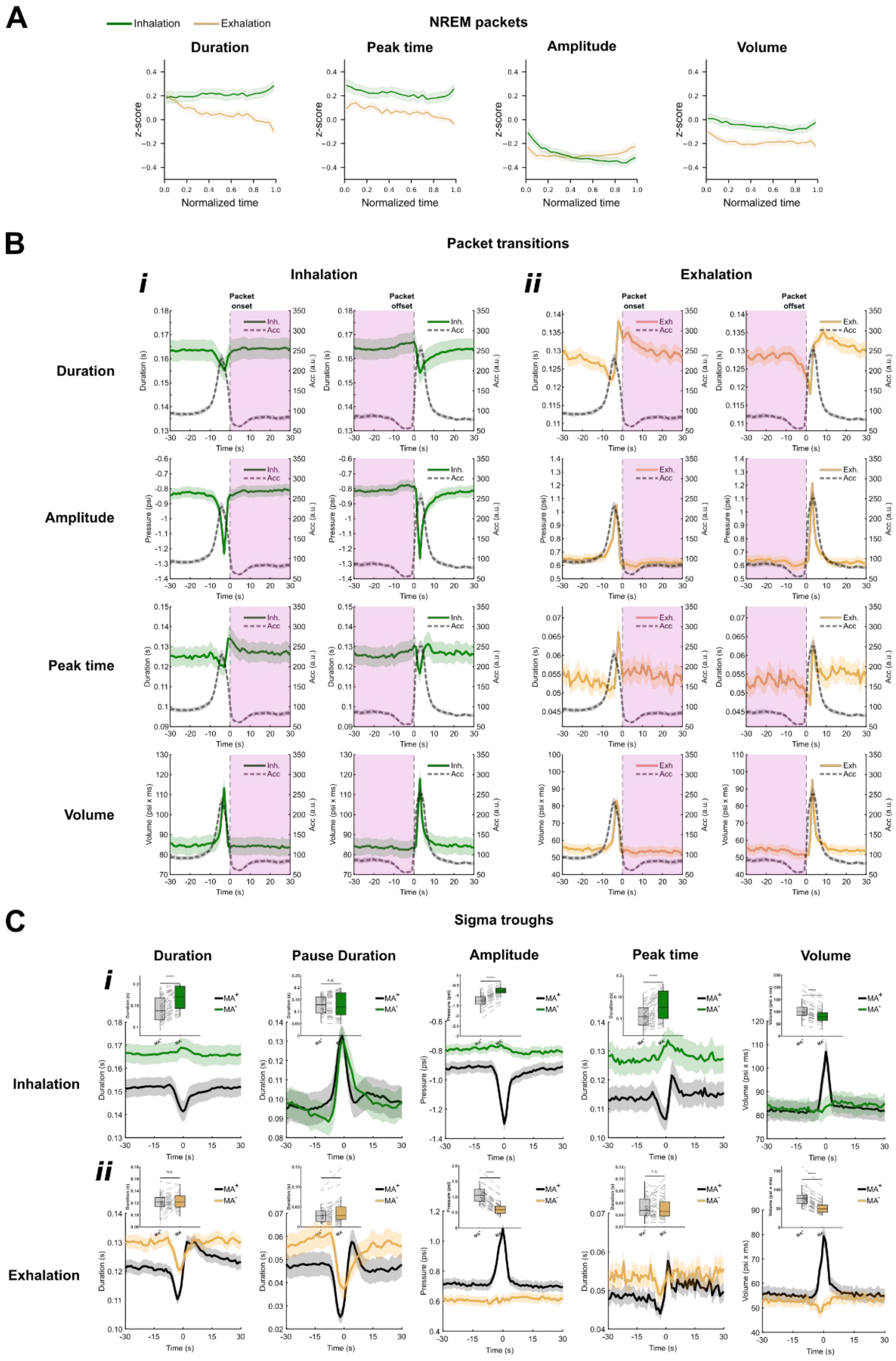

**A.** Mean (line) +/- S.E.M. (areas) curves (z-score) representing the evolution of respiratory features (duration, peak time, amplitude and volume) for inhalation (green) and exhalation (gold) during normalized time (0-1) of NREM packets. Z-scores were computed on a cycle-by-cycle basis by using only cycles in NREM for normalization (mean and SD computation).

**B.** Mean (line) +/- S.E.M. (areas) curves representing overall changes across respiratory features (rows: duration, amplitude, peak time and volume) for inhalation (***i***) and exhalation (***ii***) aligned to the onset/offset of NREM packets (light purple area) together with the mean acceleration signal (dashed grey line) representing micro-arousals.

**C.** Mean (line) +/- S.E.M. (areas) curves representing respiratory features (duration, pause duration, amplitude, peak time and volume) for inhalation (***i***) and exhalation (***ii***) aligned to sigma troughs coincident with micro-arousal ($MA^+$, black) vs without micro-arousal ($MA^-$, green for inhalation, gold for exhalation). Inlet panels show box plots representing $25^{th}$,$50^{th}$ and $75^{th}$ percentile of values at sigma troughs (t = 0) for $MA^+$ vs $MA^-$ and compared with Wilcoxon test.

***i)*** Inhalation:

a) duration mean ± SEM: $MA^+$ = 141.4 ± 4.4 ms, $MA^-$ = 169.8 ± 4.3 ms, Wilcoxon test, z = -5.32, *P* = 9.84e-8;

b) pause after inhalation duration mean ± SEM: $MA^+$ = 128.3 ± 5.6 ms, $MA^-$ = 128.4 ± 8.3 ms, Wilcoxon test, z= -0.28, *P* = 0.78;

c) amplitude mean ± SEM: $MA^+$ = -1.30.3 ± 0.05 psi, $MA^-$ = -0.78 ± 0.04 psi, Wilcoxon test, z = -5.36, *P* = 8.39e-8;

d) peak time mean ± SEM: $MA^+$ = 106.4 ± 4.0 ms, $MA^-$ = 131.6 ± 5.1 ms, Wilcoxon test, z = -5.23, *P* = 1.71e-07;

e) volume mean ± SEM: $MA^+$ = 106.9 ± 5.7 psi x ms, $MA^-$ = 83.6 ± 4.5 psi x ms, Wilcoxon test, z = 4.97, *P* = 6.80e-07.

***ii)*** Exhalation:

a) duration mean ± SEM: $MA^+$ = 120.8 ± 2.2 ms, $MA^-$ = 122.9 ± 2.0 ms, Wilcoxon test, z = -0.29, *P* = 0.77;

b) pause after exhalation duration mean ± SEM: $MA^+$ = 30.7 ± 3.1 ms, $MA^-$ = 37.3 ± 4.6 ms, Wilcoxon test, z= -2.31, *P* = 0.02;
c) amplitude mean ± SEM: $MA^+$ = 1.08 ± 0.04 psi, $MA^-$ = 0.62 ± 0.03 psi, Wilcoxon test, z= 5.36, *P* = 8.39e-08
d) peak time mean ± SEM: $MA^+$ = 52.6 ± 2.8 ms, $MA^-$ = 52.8 ± 3.0 ms, Wilcoxon test, z= 0.30, *P* = 0.77;
e) volume mean ± SEM: $MA^+$ = 79.4 ± 3.7 psi x ms, $MA^-$ = 51.3 ± 2.5 psi x ms, Wilcoxon test, z= 5.32, *P* = 1.07e-07.
